# SARS-CoV-2 shifts transcription of host gene to increase Spike acylation and boost infectivity

**DOI:** 10.1101/2023.04.15.537011

**Authors:** Francisco S. Mesquita, Laurence Abrami, Lucie Bracq, Nattawadee Panyain, Vincent Mercier, Béatrice Kunz, Audrey Chuat, Joana Carlevaro-Fita, Didier Trono, F. Gisou van der Goot

## Abstract

SARS-CoV-2 infection requires Spike protein mediating fusion between the viral and cellular membranes. The fusogenic activity of Spike requires its post-translational lipid modification by host S-acyltransferases, predominantly ZDHHC20. Previous observations indicate that SARS-CoV-2 infection augments the S-acylation of Spike when compared to transfection. Here, we find that SARS-CoV-2 infection triggers a change in the transcriptional start site of the *zddhc20* gene, both in cells and in an *in vivo* infection model, resulting in a 67-amino–acid-long N-terminally extended protein with 37-times higher Spike acylating activity, leading to enhanced viral infectivity. Furthermore, we observed the same induced transcriptional change in response to other challenges, such as chemically induced colitis, indicating that SARS-CoV-2 hijacks an existing cell damage response pathway to generate more infectious viruses.

## Main Text

Though well-studied, details of how the infectivity of ß-coronaviruses (β-CoVs), such as the causative agent of the COVID-19 pandemic, SARS-CoV-2, depend on host factors are mostly unknown. These large, enveloped RNA viruses enter host cells through interactions between the trimeric envelop glycoprotein Spike and host receptors such as ACE-2, followed by Spike-mediated fusion of the viral and cellular membranes. Like most other envelop glycoproteins (*1*), Spike undergoes multiple post-translational modifications, in particular S-acylation, which attaches a medium chain fatty acid via a thioester bond to cysteine residues in its cytoplasmic tail. This short tail contains no less than 10 such residues within a 20–amino-acid stretch (*2*),with full acylation thus resulting in 30 fatty acids per Spike trimer. Acylation has been shown to influence the lipid composition of the virions and to drastically enhance the fusogenic capacity of the produced viruses (*3–6*). The efficiency of Spike S-acylation is therefore determinant for infectivity. However, during our previous work we observed that Spike S-acylation is higher in cells infected with SARS-CoV-2 when compared to Spike transfection into the same cells (*4*).

We and others have shown that S-acylation of Spike is mediated predominantly by the ZDHHC20 acyltransferase, with a contribution from ZDHHC9 (*4, 6, 7*). Here we show that SARS-CoV-2 infection, whether in human cells or in mice, triggers the use of an upstream transcriptional start site of the *zdhhc20* gene, leading to the production of a N-terminally extended enzyme, ZDHHC20^Long^. The expression of ZDHHC20^Long^ is also triggered following chemically induced colitis and other cellular challenges, suggesting that it is part of a “damage” response pathway. ZDHHC20^Long^ drastically enhances Spike acylation, leading to the enhanced fatty acid decoration of the Spike trimers. As a consequence, the fusogenic capacity of Spike is augmented, and viral infectivity is optimized.

### Comparison of Spike S-acylation during infection vs. transfection

During our study aimed at fully characterizing Spike S-acylation, we had hints that the degree of S-acylation of the 10 available sites on SARS-CoV-2 Spike differed between infection and transfection (*4*). This is fully apparent when using PEGylation, an assay that indicates the relative abundance of acylated proteins within the population, by replacing acyl chains with 5-kDa PEG (*8*) (after hydroxylamine treatment, Methods section), which produces molecular weight shifts corresponding to the number of acylated cysteines per Spike molecule. Visualized by western blot, we found that Spike undergoes extensive S-acylation in infected cells, with a complete band shift of the Spike S2 band denoting that all Spike molecules are massively acylated following their synthesis, and that the attachment of up to 10 PEG molecules may hinder migration of fully modified Spike full-length protein in these gels (Fig. 1A) (*4*). In contrast, when a Spike-expressing vector was transfected into cells, Spike S-acylation is readily detectable by ^3^H-palmitate incorporation (*4*), yet we could not detect major band shifts upon PEGylation (Fig. 1A). This indicates that Spike S-acylation of individual Spike molecules is much less pronounced following transfection than infection.

**Figure 1:**
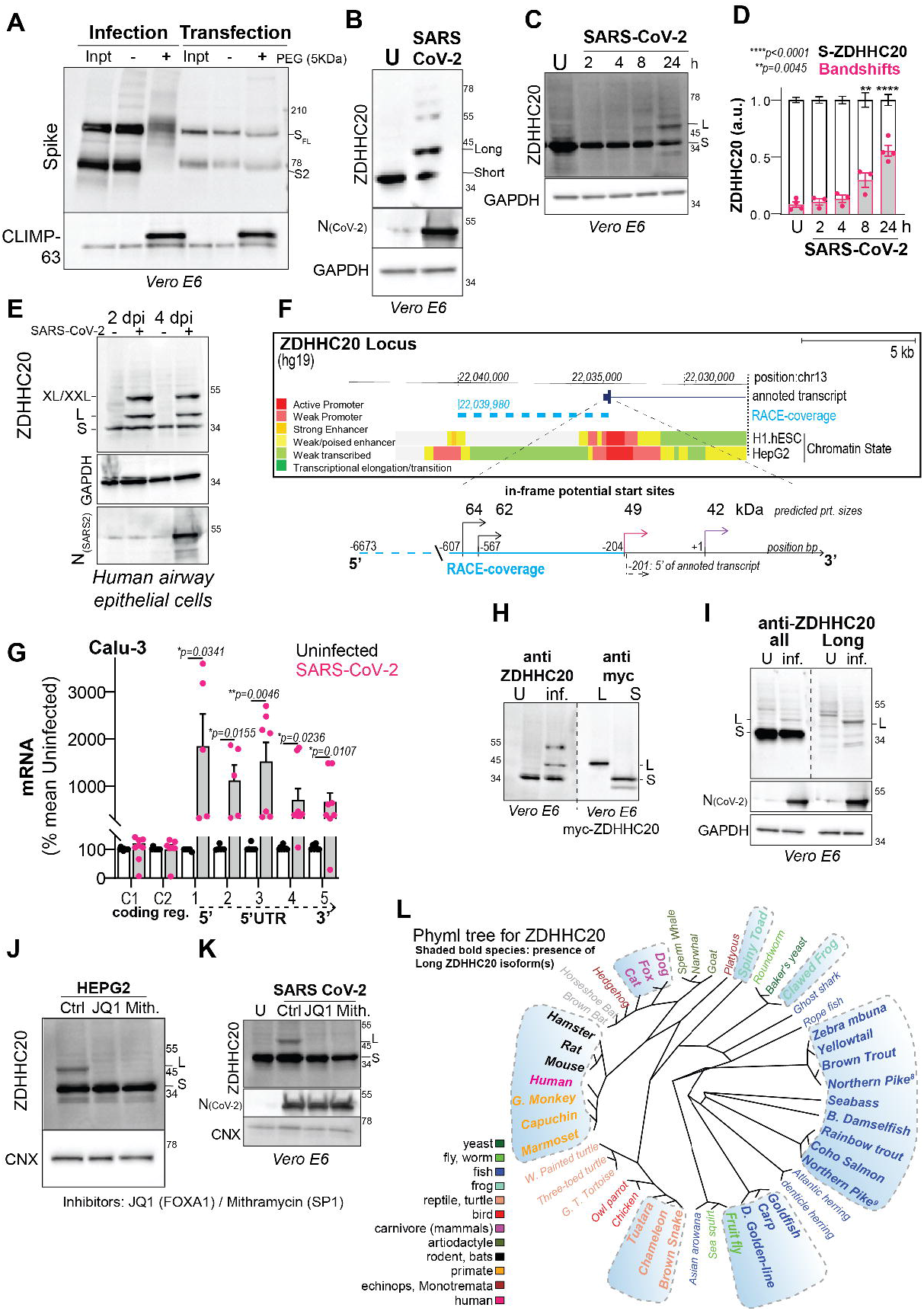
SARS-CoV-2 triggers a change in the transcriptional start site of the *zdhhc20* gene. **A.** Acyl-PEG exchange in Vero E6 cells infected with SARS-CoV-2, MOI = 0.1, 24 h (Infection) or transfected with Spike (Transfection). Western blot (WB) of input cell extract (Inpt), PEG-tagged protein (+PEG), and control (-) after hydroxylamine treatment. Spike (full length protein-SFL and cleaved fragment S2) or positive control Climp-63. **B.** WB of ZDHHC20^ALL^, nucleocapsid (N) and GAPDH (loading control) of VERO E6 cell extracts of uninfected (U) or infected as in A. Short (42 Kda, S) and Long (49 Kda, L) ZDHHC20 forms are indicated. **C.** Same as B in cell extracts infected for different times. **D.** Quantification of ZDHHC20 bandshifts in B. Results are mean ± SEM, and each dot represents one independent experiment of n = 4 (U and 24 h) and n = 3 (2, 4, 8 h). *P* values comparing shifts to U-control were obtained by two-way ANOVA, Dunnet’s multiple comparison. **E.** Same as in B in extracts of human airway epithelial, 2 and 4 days post-infection (dpi). S, L and XL/XXL (62/69 Kda) ZDHHC20 forms are indicated. **F.** Overview and zoom of ZDHHC20 locus (Human version hg19-http://genome.ucsc.edu) indicating four transcription start sites (TSS, −607, −567, −204, +1) at in-frame ATG codons (coding for XXL/XL, Long and Short respectively); 5’-RACE coverage describes the amplified genomic region; Transcript annotation corresponds to GENCODE Genes track (version V4 0lift37); Chromatin State displays segmentation data from ENCODE consortia for two different cell lines (H1.hESC and HepG2). **G.** mRNA quantification using primers probing for different locations in z*dhhc20* transcripts (coding region: C1, C2; 5’ UTR: 1–5 cover increasing lengths of 5’ ends see also fig S1D) in Calu-3 cells uninfected or infected as in A Results are mean ± SEM, and each dot represents one independent experiment. *P* comparing to uninfected, were obtained by multiple unpaired t-tests. **H.** WB of ZDHHC20 on Vero E6 extracts uninfected (U) or infected as in A (Left). WB of myc-ZDHHC20 on Vero E6 extracts transfected 24 h with ZDHHC20^Long^ (L) or ZDHHC20^Short^ (S) (Right). **I.** WB as in B (Left) compared to ZDHHC20^Long^-specific WB analysis (Right) on Vero E6 extracts uninfected (U) or infected as in B. **J.** WB of ZDHHC20 and Calnexin (loading control, CNX) on HEPG2 cell extracts treated 24 h with 100 nM JQ1 (FOXA1 inhibitor) or 100 nM mithramycin (SP1 inhibitor). **K.** WB as in B in infected Vero E6 cell extracts treated as in J. **L.** Phyml tree for ZDHHC20 in various species with presence of ZDHHC20^Long^ in shaded blue boxes.

### SARS-CoV-2 triggers a change in the transcriptional start site of *zdhhc20*

To understand how the efficiency of Spike S-acylation increases in infected versus transfected cells, we monitored the expression of the ZDHHC20 acyltransferase by western blot. In infected Vero E6 (Green monkey) cells, a cell line classically used for SARS-CoV infection assays, the enzyme migrated as multiple isoforms in addition to the 42 kDa protein also detected in control cells, with an abundant >45 kDa band and fainter ∼55 kDa bands (Fig. 1B). Such higher molecular weight species of ZDHHC20 became visible at 8 h post-infection of Vero E6 cells (Fig. 1CD) and also seen upon infection of human lung-derived Calu-3 (fig. S1A) or primary airway human epithelial cells (Fig. 1E). The appearance of higher molecular weight ZDHHC20 bands preceded significant detection of Spike or SARS nucleocapsid protein (N, Fig. 1E). This was not specific to the SARS-CoV-2 B.1-strain, since both Delta-Δ (B1.617.2) and Omicron-Ο (BA.1) triggered this same effect in ACE-2/TMPRSS2 expressing HEK293T cells (fig. S1B).

The green monkey ZDHHC20 protein sequence reported in UniProt (A0A0D9RZN5) carries an N-terminal extension that is not reported for the human protein (Q5W0Z9) (fig. S1C). We, therefore, analyzed the human *zdhhc20* gene and found 3 additional in-frame ATG sequences in the 5’ region, predicted to lead to proteins of 49, 62, and 64 kDa in addition to the 42 kDa canonical species (Fig. 1F). Since these additional putative translational start sites were all located upstream of the 5’ end of the annotated transcript, we performed 5’ Rapid Amplification of cDNA Ends (RACE) in infected cells for 5’-extended *zdhhc20* transcripts (see Methods). Some >15% of the RACE products included all 3 ATGs, spanning >6500 bp upstream of the annotated start site (Fig. 1F, fig. S1D, table S1). We next quantified *zdhhc20* mRNA by qPCR. While primers to the coding sequence led to equivalent *zdhhc20* mRNA levels in control and infected cells, primers to the 5’ extensions showed enhanced expression during infection (Fig. 1G, fig. S1DE).

To confirm that infected cells express a protein with an in-frame N-terminal extension, we generated cDNA constructs of the most abundant, 49 kDa, longer form (ZDHHC20^Long^) with an N-terminal myc-tag as well as a polyclonal antibody to peptides present in the N-terminal extension (fig. S1C, see Methods). The ZDHHC20^Long^ protein migrated similarly to the >45 kDa band expressed upon infection (Fig. 1H), and this band was recognized by the anti-peptide antibody in infected but not control cells (Fig. 1I). Thus, SARS-CoV-2 infection promotes expression of *zdhhc20* transcripts with longer 5’ regions, coding for a ZDHHC20 protein harboring N-terminal extensions, of which ZDHHC20^Long^, with a 67 amino acid extension, is the most abundant.

To understand this change in transcriptional start site (TSS), we analyzed chromatin state segmentation tracks from UCSC Genome Browser (*9*), which integrates ChIP-seq data across nine human cell types (see Methods). This showed that promoter activity was almost exclusively predicted in the region neighboring the human annotated start site, as depicted for embryonic stem cells (H1-hESC) (Fig. 1F, fig. S1F). However, in hepato-carcinoma HepG2 cells, additional promoter activity was reported approximately 6500 bps upstream from the annotated start site, proximal to the 5’ ends determined by 5’ RACE (Fig. 1F, fig. S1D-F). We probed HepG2 cells for ZDHHC20 protein expression and found constitutive expression of ZDHHC20^Long^ (Fig. 1J). To identify transcription factors (TF) binding to this upstream promotor region, we analysed ChIP-seq Peak data for HepG2 cells in ENCODE (fig. S1F). We tested the potential involvement of a panel of these TFs and found that the expression of ZDHHC20^Long^ in HepG2 cells was inhibited by siRNAs against FOXA1/2 or SP1 (fig. S1G) and by their pharmacologic inhibitors during SARS-CoV-2 infection in Vero E6 cells (Fig. 1K). FOXA1 and SP1 were also upregulated during infection in both Vero E6 and Calu-3 cells (fig S1H), but their over-expression was not sufficient to trigger ZDHHC20^Long^ production in HeLa cells (fig. S1I).

Altogether these results demonstrate that SARS-CoV-2 infection triggers the FOXA1- and SP1-mediated activation of an upstream promoter of the *zdhhc20* gene, resulting in the production of a 67–amino-acid N-terminally extended acyltransferase product. Alternative promoters are not unusual and have been found for about half of human genes (*10, 11*). The usage of alternative TSSs, recently described as a conserved stress-response mechanism, however seems to be aimed at fine-tuning mRNA expression without altering the proteome (*12*), in contrast to what we observed here. Noteworthy, genomic analyses predict the possible synthesis of N-terminally extended versions of ZDHHC20, similar to the one induced by SARS-CoV-2 infection of human cells, in a wide range of animal species, including many mammals, fishes, amphibians and the fruit fly (Fig. 1L, fig. S1C).

Interestingly, FOXA1 also coordinates the expression of the Spike receptor ACE-2 and the Spike priming factor TMPRSS2 (*13*). It remains unclear how FOXA1 and SP-1 gain access to the promoter region located 6500 base pairs upstream from the annotated start site of the human *zdhhc20* gene. This could be due to specific virus-induced alterations of host genome architecture (*14–17*). Alternatively, since the TSS of *zdhhc20* also changed in response to other cellular changes, the modification in gene architecture might be part of a more general host response to stress/infection.

### SARS-CoV-2 infection triggers the production of ZDHHC20^Long^ in mice

Before probing for the expression of ZDHHC20^Long^ in a mouse model of SARS-CoV-2 infection, we analyzed the expression of murine ZDHHC20 under physiological conditions. All tissues tested expressed exclusively ZDHHC20^Short^, except for the small intestine (Fig. 2A), where *zdhhc20*^Long^ transcripts were also detected. FOXA1 expression was also higher in the duodenum compared to kidney or colon (fig. S2A). Publicly available murine ZDHHC20 mRNA sequences (*9*) also indicate the existence of transcripts with long 5’UTRs covering alternative start sites (fig. S2B). Thus, to evaluate the distribution of *zdhhc20*^Long^ mRNAs in the duodenum, we used RNAscope probes against the extended 5’ region (fig. S2B). We confirmed the expression of a longer transcript of *zdhhc20* in the duodenum (Fig. 2B), with a higher level in the crypt region and at the base of the villi (fig. S2C). Analysis of commercially available RNA from a panel of human tissues also indicated that *zdhhc20* transcripts with longer 5’ regions were significantly increased in the small intestine (Fig. 2C). Thus, both in mouse and human, most tissues express ZDHHC20^Short^, with the exception of the small intestine, which expresses ZDHHC20^Long^.

**Figure 2:**
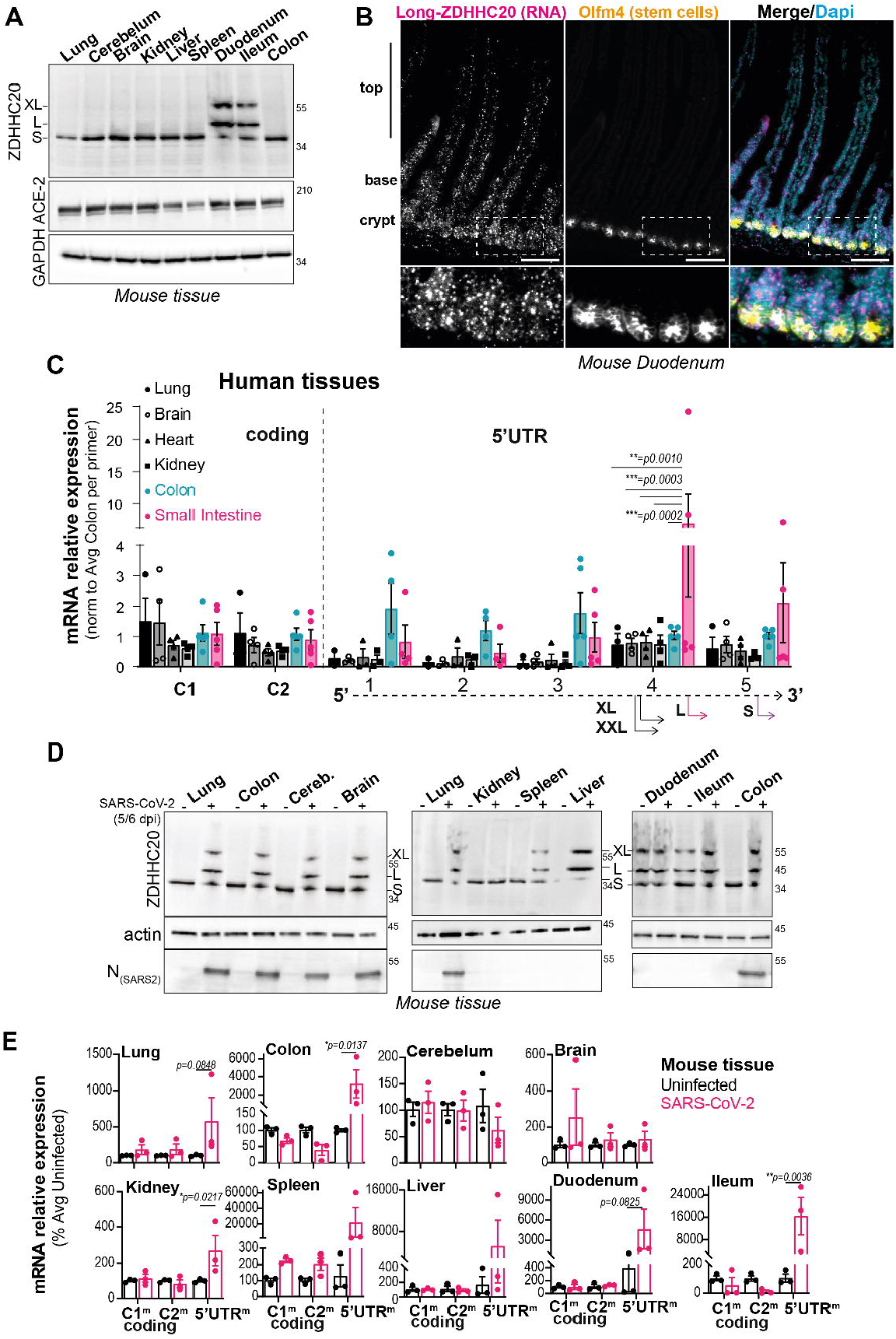
SARS-CoV-2 infection triggers expression of ZDHHC20^Long^ in mice. **A.** WB of ZDHHC20, ACE-2, and GAPDH loading control in mouse tissue extracts. Short (44 Kda, S) Long (50 Kda, L) and XL (53 Kda) ZDHHC20 forms are indicated. **B.** Longitudinal sections of mouse duodenum are labeled with an RNAscope probes for 5’ UTR *zdhhc20* RNA, DAPI, and with antibodies against Olfm4 (as marker for intestinal stem cells). Scale bars 100 µm. **C.** mRNA quantification in various human tissues, using primers probing for different locations in z*dhhc20* transcripts (coding region: C1, C2; 5’ UTR: 1–5 cover increasing lengths of 5’ ends see also fig S1D). Results are mean ± SEM and each dot represents one independent pool of human derived tissue-specific mRNA. Lung, n = 3, Brain and Heart, n = 4, and Colon and Small intestine, n = 5. *P* values comparing primer specific expression between tissues were obtained by two-way ANOVA with Tukey’s multiple comparison **D.** WB analysis of tissue extracts from control uninfected mice and mice infected intranasally with 10^4^ (plaque forming units – PFU) of SARS-CoV-2. Tissues were harvested between 5–6 days post infection and processed for WB as in A. Actin was used as loading control. **E.** mRNA quantification in different murine tissues uninfected or infected as in D, using primers for different locations in z*dhhc20* transcripts (coding region (C1, C2) or the 5’ UTR see also fig S2B). Results are mean ± SEM and each dot represent one of 4 independent mice. *P* values were obtained by two-way ANOVA with Sidak’s multiple comparison.

We next infected mice intranasally with SARS-CoV-2 and analyzed tissues for the presence of the various ZDHHC isoforms and the viral nucleocapsid N protein after 5 and 6 days. All tested tissues expressed ZDHHC20^Long^ as well as some even longer forms (XL), with the notable exception of the kidney, brain, and cerebellum (Fig. 2D). N was detected in lung, colon, brain, and cerebellum but not in kidney, spleen, liver, duodenum, and ileum (Fig. 2D). We analyzed the mRNA using primers complementary to the coding region or the 5’UTR (fig. S2B), which was consistent with the expression of 5’-extended *zdhhc20* transcripts upon SARS-CoV-2 infection in all organs except the cerebellum and brain (Fig. 2E). These observations show that under physiological conditions, ZDHHC20^Long^ is only expressed in specific mouse tissues, such as the small intestine, but becomes expressed in almost all tissues upon infection with SARS-CoV-2.

### ZDHHC20^Long^ is expressed in mouse colon following chemically induced colitis

Given the proposed association between alternative transcription and stress (*12*), and the fact that ZDHHC20^Long^ is expressed physiologically in the upper intestinal tract, we wondered whether the expression of ZDHHC20^Long^ could be triggered in the colon in response to stress. Mice were treated orally with dextran sodium sulfate (DSS), a well-established chemically induced colitis model (*18*), for 7 days and then allowed 3 days of recovery, after which we assessed *zdhhc20* expression. We detected ZDHHC20^Long^ and XL forms in the colon by western blot (Fig. 3A). RNAscope analysis showed that *zdhhc20* transcripts with long 5’UTRs were increased locally in regions of the colon undergoing apparent recovery (hypertrophic crypts with differentiated-colonocyte marker KRT20) (Fig. 3, B and C), even though the average expression in the colon was similar (fig. S3, A and B).

**Figure 3:**
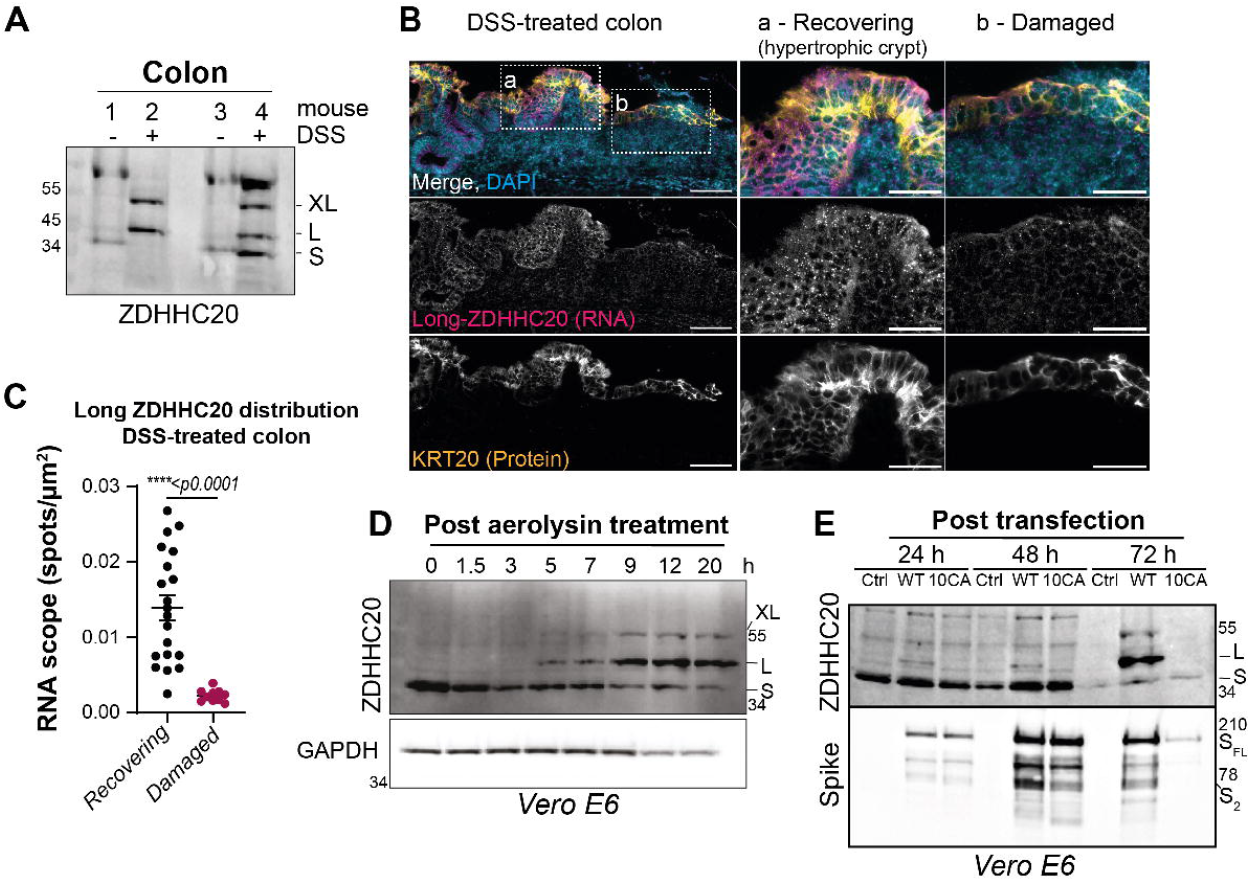
ZDHHC20^Long^ expression is triggered in mouse colon following chemically induced colitis. **A.** WB analysis of mice treated or not with DSS for 7 days followed by 3 days recovery, showing ZDHHC20 expression from colons from 2 independent mice for each condition S, L and XL forms are indicated. **B.** RNAscope analysis of DSS-treated mice, as in A, using probes for 5’ UTR *zdhhc20* RNA (see fig. S1B), DAPI staining, and anti-KRT20 antibodies (to mark intestinal epithelial cells). Scale bars 100 μm. **C.** Representative quantification (one of 3 independent mice) of Long-*zdhhc20* RNA scope spots per µm^2^ in colon sections from mice DSS-treated as in A. Results are mean ± SEM and each dot corresponds to one field-of-view selected from recovering (n=20) or damaged (n=10) zones represented in B. *P* values were obtained by unpaired t test. **D. E.** WB analysis of ZDHHC20 expression (S, L and XL forms are indicated) in Vero E6 cells treated D 1 h with 10 ng/ml of proaerolysin at 37°C, washed, and further incubated at 37°C for the indicated times; or **E.** transfected or not for 24, 48 or 72 h with WT or 10CA Spike. GAPDH shows loading control in D and Spike levels were monitored in E.

As our results suggest that ZDHHC20^Long^ expression could constitute a general response to danger, we tested yet another challenge, treating cells with the pore-forming toxin aerolysin, which leads to loss of plasma membrane integrity (*19*). We chose a transient exposure to aerolysin (1 h), which triggers cell-survival signaling (*20*). In toxin-exposed Vero E6 (Figs. 3D) and HeLa cells (fig S3C), ZDHHC20^Long^ expression was visible within 5 h and became increasingly abundant over time; the expression disappeared after 48 h (Fig. S3D).

Finally, we noticed that transfecting cells with wildtype (WT) Spike, but not its S– acylation-deficient mutant (10CA, all cytosolic cysteines mutated (*4*)), also triggered mild ZDHHC20^Long^ expression (Fig. 3E). Since Spike induces cell-to-cell fusion in an acylation-dependent manner (*5, 21*), this again could be the expression of ZDHHC20^Long^ in response to danger.

### ZDHHC20^Long^ is longer lived and retained in the ER

We next investigated the consequences of the N-terminal 67–amino-acid extension. Metabolic labelling with ^35^S Cys/Met showed that transfection of cells with equivalent amounts of the corresponding plasmid DNAs led to the synthesis of the same amounts of Long or Short ZDHHC20 (fig. S4A). However, total ZDHHC20^Long^ levels were approximatively 5 times higher than ZDHHC20^Short^ (Fig. 4A), which could be explained by a 2-fold increase in the apparent protein half-life, determined by ^35^S Cys/Met metabolic pulse-chase analysis (Fig. 4B).

**Figure 4:**
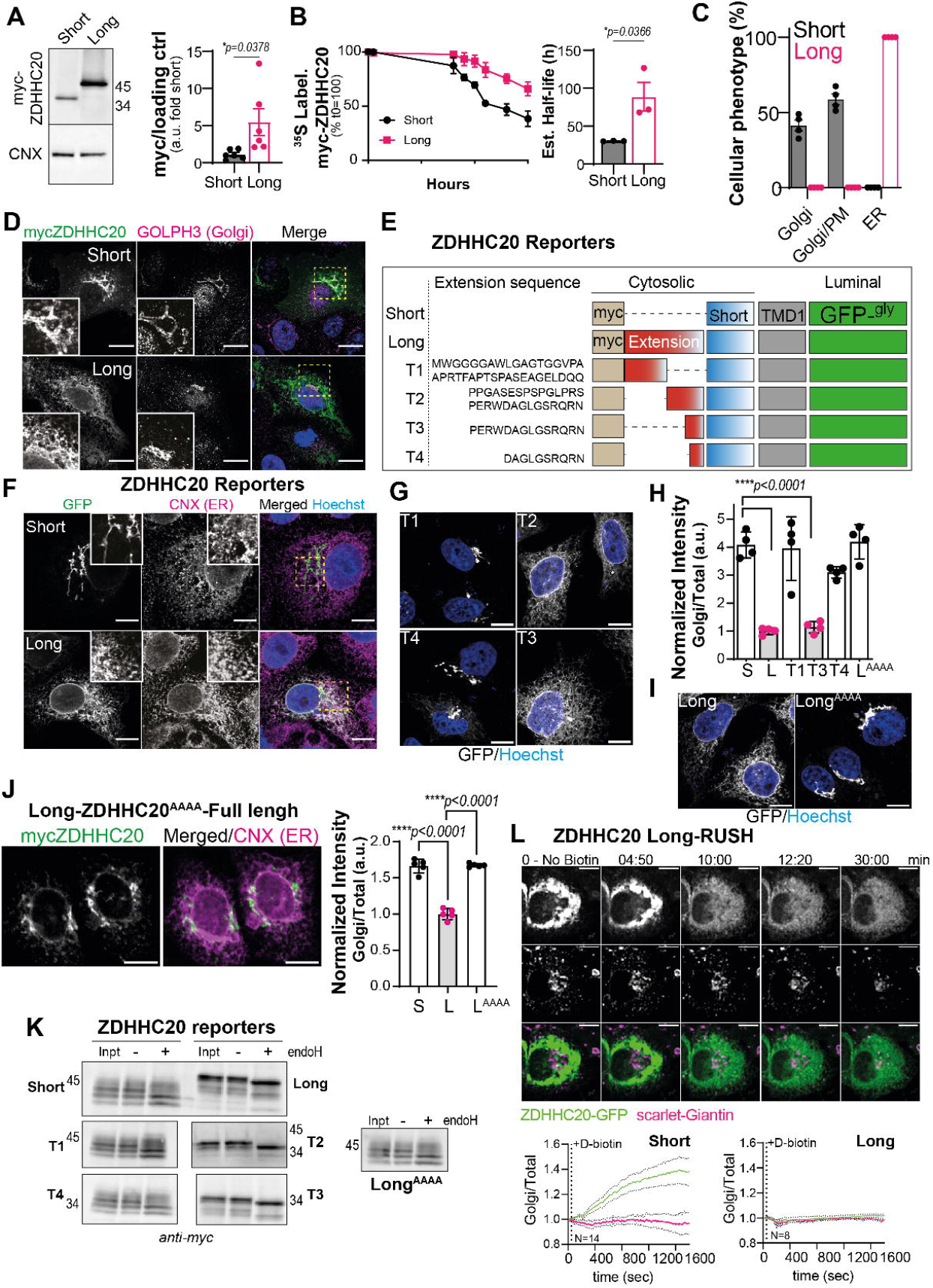
The N-terminal extension of ZDHHC20^Long^ controls its abundance and localization. **A.** WB of myc-ZDHHC20 and calnexin (CNX) on Vero E6 cell extracts transfected 24 h with ZDHHC20^Short^ (Short) or ZDHHC20^Long^ (Long). Ratio of myc-ZDHHC20 to loading control corresponds to mean + SEM, and each dot represents one of n = 6 independent experiments. **B.** Hela cells expressing myc-ZDHHC20 (Short or Long) metabolically labelled with 35S-Met/Cys for 20 min and chased as indicated. 35S-Met/Cys apparent decay in ZDHHC20 immunoprecipitation fractions normalized to T = 0, corresponds to mean ± SD, of n = 3 independent experiments. Estimated half-lives were extracted from the individual experiments using non-linear regression with one-phase decay. Results are meanLJ±LJSEM and *P* values obtained by unpaired t test. **C.** Qualitative quantification of myc-ZDHHC20 (Short or Long) distribution in Golgi, Golgi and plasma membrane (PM), or ER in Vero E6 cells. Results are mean ± SEM, of n = 4 independent experiments where a total of 1235 (Short) and 814 (Long) cells were counted. **D.** Immunofluorescence (IF) of Vero E6 cells expressing myc-ZDHHC20 (Short or Long) labelled for myc (ZDHHC20) and Golgi marker GOLPH3, scale bar: 10 µm. **E.** Illustration of different ZDHHC20 reporters containing a myc-tagged short ZDHHC20 N-terminal tail, the first transmembrane (TM) domain, and a luminal glycosylated GFP. The indicated N-terminal extensions were added for each reporter. **F.** IF of Vero E6 cells expressing ZDHHC20^Short^- or ZDHHC20^Long^-reporters labelled for ER marker calnexin (CNX) and nuclear-stained with Hoechst, scale bar: 10 µm **G.** IF of HeLa cells expressing the indicated ZDHHC20-reporters illustrated in E, scale bar: 10 µm. **H.** Quantification by high-throughput automated microscopy of Hela cells expressing the indicated ZDHHC20-reporters for 24 h. The data represent ratio of GFP signal between Golgi and cytoplasm normalized by to ZDHHC20^Long^-reporter. Data, representative of other 2 independent experiments, were obtained from 4 wells with 9 images per well. Each dot represents the mean from one independent well, results are mean ± SEM and *P* values obtained by one-way ANOVA with Tukey’s multiple comparison. The total number of cells analyzed was respectively: S:3755, L:3096, T1:2990, T2:3213, T3: 3078, T4: 3681. **I.** IF of HeLa cells expressing ZDHHC20^Long^-reporter with PERW/AAAA mutation, scale bar: 10 µm. **J.** IF of HeLa cells expressing full length ZDHHC20^Long^ with PERW/AAAA mutation (L^AAAA^) labelled for myc and CNX, scale bar: 10 µm. High-throughput quantification, as in H, in Hela cells expressing the indicated myc-tagged ZDHHC20 reporters. Data were obtained from 5 wells with 25 images per well. Each dot represents the mean from one independent well, results are mean ± SEM and *P* values obtained by one-way ANOVA with Tukey’s multiple comparison, results are mean ± SEM. The total number of cells analyzed was respectively: S:19478, L:2041 and L^AAAA^: 19166. **K.** EndoH assay: WB on Vero E6 cell extracts expressing the indicated ZDHHC20- reporters. Cell extracts (40 µg) were treated (+) or not (-) with EndoH **L.** Time-lapse confocal microscopy images of Vero E6 cells co-expressing (24 h) the Golgi marker Scarlet-Giantin, ZDHHC20^Long^-RUSH-GFP reporter and an ER-resident hook. Synchronized trafficking was monitored upon D-biotin addition, after T_0_. The normalized GFP ratio at the Golgi (Scarlet-Giantin) was calculated throughout time (T_0_=1; every 10 sec for 30 min) for n = 14 ZDHHC20^Short^- or n = 8 ZDHHC20^long^- RUSH-reporter expressing independent cells (see also fig S4F and movies S1-2).

We next analyzed the subcellular localization of ZDHHC20^Long^, knowing that the canonical human ZDHHC20 (ZDHHC20^Short^) accumulates in the Golgi, in vesicles, and at the plasma membrane (*4, 22*) (Fig. 4, C and D). Surprisingly, ZDHHC20^Long^ (human or monkey) localized exclusively to the ER (Fig. 4, C and D, fig. S4, B–D). To determine which part of the N-terminal extension was responsible for this drastic change in localization, we generated reporter constructs encoding chimeric proteins with the Short ZDHHC20 cytosolic tail, the first transmembrane domain of the protein fused to a GFP variant harboring a N-glycosylation site on the luminal side (Fig. 4E). On the cytosolic side, we added or not full-length or truncated versions of the N-terminal extension of ZDHHC20^Long^ (Fig. 4E). Short and Long reporters behaved in accordance to the full-length ZDHHC20 proteins, but with a more exclusive Golgi distribution for the Short reporter (Fig. 4F, fig. S4E). The T1 truncation harboring the N-terminal half of the 67–amino-acid extension also localized entirely to the Golgi, while the T2 and T3 truncations carrying C-terminal parts of the extension localized to the ER (Fig. 4, G and H). We further truncated T3 by 4 residues, namely PERW (T4), and this abolished ER localization. Mutation of these 4 residues to alanine in the Long reporter was sufficient to shift its localization from ER to Golgi (Fig. 4, H and I). Quantitative high throughput microscopy showed that a full-length AAAA ZDHHC20^Long^ mutant similarly shifted its localization from ER to Golgi, confirming the requirement for the PERW motif (Fig. 4J). These morphological observations were confirmed biochemically using Endoglycosidase-H (EndoH), which cleaves simple but not complex N-linked sugars generated in the Golgi (*23*). Characteristic of complex Golgi-generated sugars, the Short reporter as well as T1 and T4 truncations and the Long-AAAA mutant, in part migrated as a higher molecular weight smear of multiple bands, which were insensitive to EndoH treatment (Fig. 4K). In contrast, the Long reporter, and the T2 and T3 truncations did not show the higher molecular weight smear, and all protein bands were sensitive to EndoH treatment (Fig. 4K), indicating that these forms had not reached the Golgi.

To test whether the N-terminal extension leads to ER retention or ER retrieval, we used the RUSH system, wherein ZDHHC20 reporters were hooked to the ER via a streptavidin-binding peptide (SBP) and the co-expression of a streptavidin-tagged ER “hook” protein (see Methods) (*24*). After the addition of biotin, which releases the SBP-containing ZDHHC20 reporter from the hook, we monitored its synchronized transport by live microscopy. The ZDHHC20^Short^-RUSH reporter was transported from the ER to the Golgi (Fig. 4L bottom left quantification panel, fig. S4F, movie S1), whereas the ZDHHC20^Long^-RUSH reporter remained in the ER even after 30 min (Fig. 4L, movie S2). Altogether these observations show that ZDHHC20^Long^ accumulates in the ER via a specific retention motif, and has an extended half-life that leads to higher protein levels compared to ZDHHC20^Short^.

### ZDHHC20^Long^ has greatly enhanced performance in modifying Spike

Having established that the N-terminal extension present in ZDHHC20^Long^ leads to higher proteins levels than ZDHHC20^Short^ and to ER localization, we assessed its ability to S-acylate Spike, by monitoring incorporation of ^3^H-palmitate. Palmitate incorporation was observed with both forms of the enzyme. However, in cells expressing ZDHHC20^Long^, ^3^H-palmitate incorporation into Spike was much faster and also plateaued at a level ≈37 times higher than in cells expressing ZDHHC20^Short^ (Fig. 5A). This shows that in the presence of ZDHHC20^Long^, the Spike population acquires palmitate on more cysteines. Our previous mutagenesis analysis indicates that when acylation starts on a given Spike molecule, it tends to proceed to near completion on the 10 cysteines of the cytosolic domain (**<u>CC</u>**XXX**<u>CC</u>**X**<u>C</u>**XXX**<u>CC</u>**X**<u>C</u>**XX**<u>CC</u>**) (*4*). This is consistent with our previous work showing that the presence of successive cysteines leads to cooperative S-acylation (*25*), presumably because when cysteines are sufficiently close, acyl transferases can modify more than one during an enzyme-substrate contact event. The increased ^3^H-palmitate incorporation we observed in the presence of ZDHHC20^Long^, therefore suggests that more Spike molecules undergo acylation. Our previous observations also indicate that acylation of successive cysteines protects proteins from deacylation (*25*). This altogether would predict that the faster Spike gets acylated, such as in ZDHHC20^Long^-expressing cells, the more resistant it becomes to deacylation. To test this, we conducted pulse-chase experiments following ^3^H-palmitate labelling. In cells expressing ZDHHC20^Short^, Spike lost ≈78% of the ^3^H-palmitate within 6 h (Fig. 5B), as observed previously (*4*). In cells expressing ZDHHC20^Long^, only ≈33% of the Spike-associated ^3^H-palmitate was lost (Fig. 5B). Spike deacylation is thus significantly reduced in cells expressing ZDHHC20^Long^.

**Figure 5:**
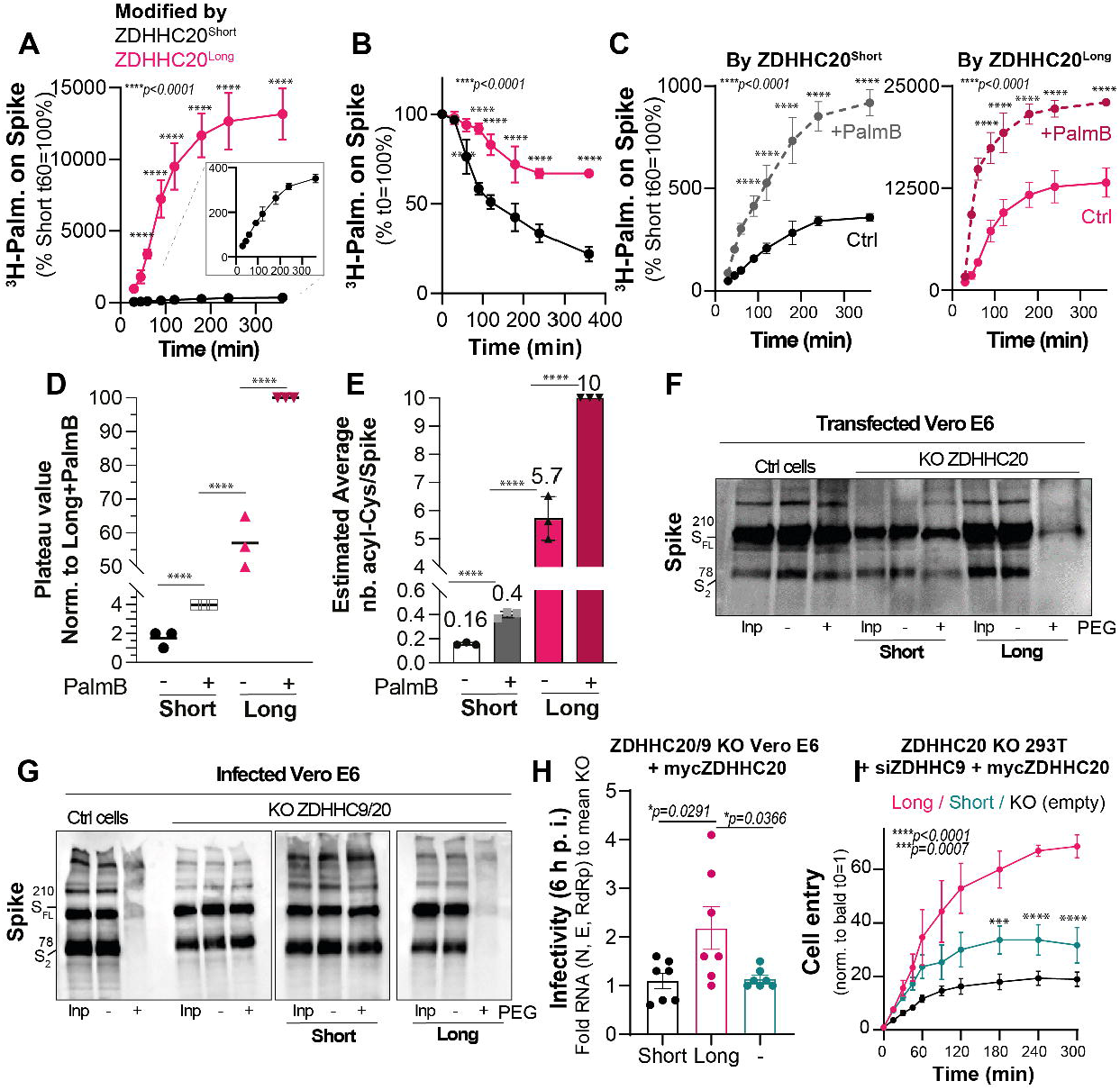
Expression of ZDHHC20^Long^ drastically increases Spike S-acylation and leads to more infectious viruses. **A.** Incorporation of ^3^H-palmitic acid in Spike-HA immunoprecipitation fractions normalized to T = 60 min from Vero E6, KO-ZDHHC20 recomplemented with ZDHHC20^Short^ or ZDHHC20^Long^. Inset highlights curve from ZDHHC20^short^-expressing cells. **B.** Quantification of ^3^H-palmitate turnover from Spike-HA immunoprecipitation fractions. Vero E6, as in A, were metabolically for 3 h pulse and chase in complete medium for indicated times. Values for each condition are set to 100% at T = 0. **C.** Spike-HA incorporated ^3^H-palmitic acid, as in A, in cells are pretreated or not (Ctrl) with thioesterase inhibitor Palmostatin B. For **A-C** results are mean ± SD of n = 3 independent experiments and *P* values were obtained by two-way ANOVA with Sidak’s multiple comparison. **D.** Average plateau values from curves depicted in C normalized to 100%-modified for cells expressing ZDHHC20^Long^ and treated with Palmostatin. **E.** Estimated number of palmitoylated Cysteine residues in Spike retrieved from plateau levels in D normalized to full 10 palmitoylated Cysteines per Spike for cells expressing ZDHHC20^Long^ and treated with Palmostatin. Average number of modified residues are indicated. For **D and E,** *P* values were obtained by one-way ANOVA with Tukey’s multiple comparison **F.** Acyl-PEG exchange in Vero E6 control (Ctrl) or KO ZDHHC20 recomplemented with ZDHHC20^Short^ or ZDHHC20^Long^ transfected with WT-Spike for 24 h. WB of Spike showing input cell extract (Inp), mass-tagged protein (+PEG) and control (-mPEG) after hydroxylamine treatment (NH_4_OH). **G.** Same as in **(F)** in Vero E6 KO-ZDHHC9/20 recomplemented with ZDHHC20^Short^ or ZDHHC20^Long^ and infected with SARS-CoV-2, MOI = 0.1, 24 h. **H.** Virion supernatants from Vero E6 KO-ZDHHC9/20 recomplemented with ZDHHC20^Short^ or ZDHHC20^Long^ and infected as F were harvested between 24-48 h post inoculation (p.i.) and adjusted to comparable viral (N and E) RNA content. Adjusted supernatants were used to re-infect naive Vero E6 and infectivity assessed by quantifying viral (N, E, RdRp) RNA 6 hours p.i.. Results normalized to mean infectivity of KOZDHHC20/9- derived supernatants are mean ± SEM, of n = 7 independent experiments. *P* values were obtained by one-way ANOVA with Tukey’s multiple comparison. **I.** Representative viral-like particles-(VLP)-Cell entry assay. Stock suspensions of reporter VLPs (produced in KO-ZDHHC20 HEK293T cells transfected with siZDHHC9 for 72 h) stably expressing ZDHHC20 (Short or Long) or empty plasmids (KO empty) were adjusted by HiBit-N input multiplicities and used for synchronized cell entry assays in ACE2-LgBit- transfected HEK293T-ACE2-TMPRSS2 target cells. Results are mean ± SEM of one representative experiment with 3 stock replicates repeated for 3 independent times. P values were obtained by two-way ANOVA with Sidak’s multiple comparison.

However, as the deacylation of Spike still occurs, to monitor the full acylation capacity, we performed ^3^H-palmitate labelling in the presence of the broad de-acylation inhibitor Palmostatin B (PalmB). Comparison between the steady state ^3^H-palmitate incorporation values can provide information on the relative number of S-acylated cysteines under the different conditions. The plateau values of ^3^H-palmitate incorporation into Spike significantly increased both in cells expressing ZDHHC20^Short^ and ZDHHC20^Long^ in the presence of Palmostatin B (Fig. 5, C and D). We assumed that the ^3^H-palmitate plateau value reached in Palmostatin–B-treated cells expressing ZDHHC20^Long^ corresponded to 100% of Spike molecules modified on all 10 cysteines (Fig. 5E). With this assumption, Spike would be modified, on average, on: (i) 5.7 cysteines in cells expressing ZDHHC20^Long^ in the absence of Palmostatin B; (ii) 0.16 cysteines in cells expressing ZDHHC20^Short^; and (ii) 0.4 cysteines in the same cells treated with Palmostatin B (Fig. 5E).

These estimations reveal that ZDHHC20^Short^ is extremely inefficient in acylating Spike. We confirmed this by PEGylating Spike from cells expressing exclusively ZDHHC20^Short^, wherein no significant band shift was observed (Fig. 5F). Even when infecting double ZDHHC9/20 KO cells complemented with ZDHHC20^Short^, PEGylation did not shift the Spike bands (Fig. 5G). In contrast, upon either transfection or infection of cells expressing ZDHHC20^Long^, Spike underwent massive PEGylation leading to almost full disappearance of the full length and S2 Spike bands (Fig. 5, F and G). Altogether, these observations show that ZDHHC20^Long^ is a drastically more potent enzyme for S-acylation of Spike. This is not only due to the increased expression of ZDHHC20^Long^, since when adjusting the amount of transfected plasmid DNA to reach similar protein levels (fig. S5A), ZDHHC20^Long^ was still far more efficient in S-acylating Spike (Fig. 5A).

ER localization might make ZDHHC20^Long^ more adapted to Spike, as SARS-CoV-2 virus assembly occurs in the ER-Golgi intermediate compartment (ERGIC) (*26, 27*). Spike can reach the ERGIC either by anterograde trafficking from the ER or retrograde trafficking from the Golgi (*28*). In cells expressing ZDHHC20^Short^, located in the Golgi, Spike would require transport to the Golgi to become acylated followed by subsequent retrograde trafficking to the ERGIC for incorporation into virions. In contrast, ZDHHC20^Long^ expression would enable Spike acylation in the ER and anterograde delivery to the ERGIC for incorporation into virions. At this stage, though, we cannot exclude that the N-terminal extension additionally improves the enzymatic activity and/or substrate recognition of ZDHHC20^Long^.

Spike S-acylation upon transfection into culture cells has been monitored by many laboratories (*3, 4, 6, 7, 29, 30*), yet we all overlooked the fact that ZDHHC20^Short^, which we now know was the main enzyme expressed in these cells, is very inefficient in modifying Spike. This is likely because ^3^H-palmitic acid incorporation and acyl-capture assays monitor gains of signal such that abundant proteins with numerous cysteines, like over-expressed Spike, will produce readily detectable signals suggestive of extensive acylation. However, these methods do not inform on the stoichiometry of S-acylation. PEGylation experiments have been done, but the minimal band shifts of Spike that were observed were not interpreted (*4, 7, 29, 31*), yet did still indicate that acylation under these conditions was occurring on very few Spike molecules within the population. Instead, Spike only becomes extensively acylated during infection when cells express significant levels of ZDHHC20^Long^.

### ZDHHC20^Long^ leads to more infectious SARS-CoV-2 viruses and viral-like particles

We next determined the consequence of the differential acylation of Spike on infectivity. To identify the specific contributions of ZDHHC20^Short^ and ZDHHC20^Long^, we expressed these enzymes in Vero E6 cells after KO of endogenous ZDHHC20 and KD of ZDHHC9 by RNAi. Following infection with SARS-CoV-2, the supernatants containing viruses were collected at 24 and 48 h. The relative abundance of structural SARS-CoV2 proteins, Spike, N and M was not significantly influenced by which acyltransferase was expressed (fig. S5B) (*4*). Following normalization to their viral RNA content, these supernatants were used to infect Vero E6 cells. Viral RNA was measured 6 h post inoculation, corresponding to a single round of infection. Infection with virions produced by ZDHHC20^Long^ expression cells induced an approximately 2-fold increase in viral RNA, indicating their higher infectivity (Fig. 5H).

Since ZDHHC20 modifies other SARS-CoV2 proteins (*32*) as well as antiviral proteins (*33, 34*), we more specifically addressed the role of ZDHHC20-modified Spike using the viral-like particles (VLPs) established by the Gallagher lab (*4, 35, 36*). These VLPs, based on a split reporter system engineered into the N protein of SARS-CoV-2, were produced from ZDHHC20 KO cells silenced for ZDHHC9 expression and re-complemented with ZDHHC20^Short^ or ZDHHC20^Long^. VLPs were normalized to their N-reporter content, and fusion dynamics were monitored by measuring bioluminescence as a function of time. VLPs displayed some fusogenic activity in the absence of Spike acyltransferases, which was moderately enhanced upon ZDHHC20^Short^ expression (Fig. 5I) as observed previously using HIV-based viral pseudotypes (*4*). In contrast, a >2-fold increase was observed when expressing ZDHHC20^Long^ (Fig. 5, H and I). We conclude that the ZDHHC20 enzyme isoform that is expressed when cells are infected by SARS-CoV-2 ensures that Spike is extensively S-acylated to optimize its fusogenic capacity.

### Likely effects of this cellular response pathway on SARS-CoV-2 infection

Thus, overall, the change in the *zdhhc20* TSS led to longer transcripts carrying 1–3 additional in-frame translational start sites, enabling the expression of ZDHHC20 proteins with extended N-termini, the most abundant of which contain an additional 67 amino acids. This extension lengthens the half-life of the protein, leading to 5-fold increase in abundance. It also contains a 4–amino-acid ER retention motif, PERW, that prevents transport to the Golgi and plasma membrane, where ZDHHC20 normally localizes. These novel characteristics contribute to the highly enhanced capacity of ZDHHC20^Long^ to modify Spike, which can then exit the ER as an essentially fully acylated trimer, ready for incorporation into newly forming virions in the ERGIC compartment. Infection thus triggers a transcription change that is exploited by SARS-CoV-2 to form more infectious viral particles. Since ZDHHC20 also modifies hemagglutinin, the envelop protein of Influenza A virus (*33*), it will be interesting to determine whether a similar change of enzyme operates during flu infection.

During SARS-CoV-2 infection, cells require ≈24 hrs or more to acquire a significant population of ZDHHC20^Long^ (Fig. 1D). During the first rounds of virus production, 6–12 hrs post infection, the produced virus will thus harbor mainly non-acylated Spike proteins. What must follow is a period of extreme heterogeneity in Spike acylation, ranging from non-acylated trimers to trimers carrying 30 acyl chains. This heterogeneity will influence the size, lipid composition and infectivity of the virions (*4, 5*). How the low infectivity of viruses produced early during infection influence the onset of COVID19 symptoms and virus spreading remains to be investigated.

Our study opens numerous questions regarding Spike. For example, does acylation of a Spike trimer enhance its probability of being incorporated into a nascent virus? Why do enveloped viruses carry so many Spike trimers? Is this to ensure that at least some are fully competent for binding and fusion? Does full Spike trimer S-acylation affect the orientation/flexibility of Spike at the surface of the virus, thus influencing its binding capacity? Finally, since ZDHHC20^Long^ is expressed in the small intestine, does infection at this site contribute to accelerated infection due to the immediate production of optimized viruses? While SARS-CoV-2 has highjacked ZDHHC20^Long^ expression to its own advantage, the expression of this modified enzyme also occurs in response to loss of plasma membrane integrity or chemically induced colitis, raising the possibility that it is part of a cellular repair pathway. Further dissection of this pathway and of the corresponding damage sensing mechanism holds the promise of interesting findings.

## AKNOWLEGMENTS

We thank C. Iseli and N. Guex for ZDHHC20 Phyml tree realization.; BIOP EPFL Core Facility; S. Vossio and D. Moreau from ACCESS Geneva; P. Turelli, EPFL for the different viral strain production; F. Perez and G. Boncopain for RUSH plasmid constructs; Machamer laboratory for E and M. antibodies (Cohen et al., 2011); all G. Van der Goot lab and D. Trono lab members for discussions. This work was supported by the Swiss Nationnal Science Foundation (SNSF-31CA30_196651) and by CARIGEST S.A.

## AUTHOR CONTRIBUTIONS

Conceptualization, F.S.M., L.A., and F.G.v.d.G.; investigation, F.S.M., L.A., B.K., L.B., N.P., V.M.., A.C., and J.C-F; funding acquisition, F.S.M., and F.G.v.d.G.; writing – original draft, F.S.M., L.A., and F.G.v.d.G.; writing – review & editing, F.S.M., L.A., L.B.,V.M.,J.C-F D.T., and F.G.v.d.G.; resources, B.K., A.C., and L.A..

