## Supplementary Materials for "SARS-CoV-2 shifts transcription of host gene to increase Spike acylation and boost infectivity"

**The PDF file includes:**

Materials and Methods
Figs. S1 to S5
Tables S1

**Other Supplementary Materials for this manuscript include the following:**

Movies S1 to S2

### Materials and Methods

#### Antibodies

ACE-2 (Abcam: ab15348; RRID: AB_301861; rabbit: 1:2000 dilution).
Actin (Millipore: MAB1501: RRID: AB_2223041; mouse: 1:4000 dilution).
Calnexin (Millipore: MAB3126: RRID: 2069152; mouse: 1:2000 dilution).
Climp63/CKAP4 (Bethyl Laboratories: A302-257A; RRID: AB_1731083; rabbit: 1: 2000 dilution).
Flag (Sigma: F3165; RRID: AB_259529; mouse : 1:2000).
GAPDH (Thermofisher: 398600; RRID: AB_2533438; mouse: 1:4000 dilution).
Giantin (Abcam: ab37266; RRID: AB_880195; rabbit: 1:200 dilution).
GOLPH3 (Abcam: ab98023; RRID:AB_10860828; rabbit : 1:200 dilution).
GM130 (BD:610823; RRID: AB_3998141; mouse: 1:200 dilution).
HA (Roche: 11867423001; RRID: AB_390918; rat: 1:500 dilution).
Keratin20 KRT20 (Cell Signalling: 13063; RRID: AB_2798106; rabbit: 1:600 dilution).
myc (Sigma: M4439; RRID: AB_439694; mouse: 1:2000 dilution).
Nucleocapside N SARS-CoV-2 (Genetex: GTX135357; RRID: AB_2868464; rabbit 1:2000 dilution).
SARS-CoV-1/2 E and M antibodies are gift from Machamer lab.
OLFM4 (Cell Signalling: 39141; RRID:AB_2650511; rabbit: 1:250 dilution).
Spike SARS-CoV-2 (Lifespan: LS-C19510; RRID: AB_840148; rabbit: 1:2000 dilution).
ZDHHC20 $^{ALL}$ (Sigma: SAB4501054; RRID: AB_10744838; rabbit: 1: 2000 dilution).
Mouse-HRP (GEHealthcare: NA931V; RRID: AB_772210; mouse: 1: 3000 dilution).
Rabbit-HRP (GEHealthcare: NA934V; RRID: AB_772206; rabbit: 1: 3000 dilution).
Mouse-Alexa488 (ThermoFisher Scientific: A-11029; RRID: AB_2534088; 1:800 dilution).
Mouse-Alexa568 (ThermoFisher Scientific: A-11037; RRID: AB_2534013; 1:800 dilution).
Rabbit-Alexa488 (ThermoFisher Scientific: A-21206; RRID: AB_2535792; 1:800 dilution).
Rabbit-Alexa568 (ThermoFisher Scientific: A-11042; RRID: AB_2534017; 1:800 dilution).
DAPI (ThermoFisher Scientific: D1306; RRID: AB_2629482; 1:5000 dilution).
Hoechst (Sigma: 94403; 1:5000 dilution).

#### Polyclonal Antibodies production against Long ZDHHC20 $^{Long}$

The 2 following peptides were used in combination to produce polyclonal antibodies in 2 different rabbits, 2 serum were pooled and were immunopurified against the 2 peptides (Peptide 1: C-EAGELDQQPPGASES-coNH2- Peptide 2: C-LPRSPERWDAGLGSRQR-coNH2).

#### Compounds and reagents

D-Biotin (Combi-Blocks: CSLSS-7910)
ENDO H (Bioconcept: P0702L)
Hydroxylamine (Sigma: 55460)
JQ1(Sigma: SML-1524; used final concentration 100nM, FOXA1 inhibitor)
Mithramycin (LKT: LKT-M3476; used final concentration 100nM, SP1 inhibitor)
NEM (Sigma: E1271)
Palmostatin B (Calbiochem: 178501; used final concentration 50□M, thioesterase inhibitor)
PEG-5KDa (Sigma: 63187)
TCEP (Sigma: C4706)
Zebra-spin desalting columns (Pierce: PIER89882)
3H-palmitic acid (American Radio-labelled Chemicals: ART0129-25)
35S-met-cys (Hartmann Analytic GmbH: IS103-185)

**RNAscope probes for *in situ* hybridization**
A mix of 7ZZ Probes tagged C3 were designed and produced by Advanced cell diagnostic (ACD) in the following mouse ZDHHC20 sequence:
5’-GAGTCTTATATTTAAGTATATATAATATTTTTCTTGCTTGCACGCTTGAG
TGATGCACAGCCTTATGTAATAGGGAACCCGACTTGTATGGGTCCTGAAA
GACCTAGGGGAAGAGTGTTATACAGAGTGCCACCCAAGGCACACAGGGCA
GCTAAGAATTCATTCCTCTCCCCTTGCGCATCCCCATCCCTCTCGTCACC
CAGTCCTAGATGGCCTCCTACACATCCTTAGCACTCTCCTCTCTTTTCCA
CCAAGGCACCCCCAAATCCCATGGCGGGCCTGAGCAGAAGCCCCGCCCC
AACTTCAGGCCCCGCCTCCTTCGGCCGGGTAGCCCCACCCCCTACGGGGA
TTGCCAGGCGCGGGAC-3’

RNAscope Multiplex Fluorescent V2 assay was performed according to manufacturer's protocol 4 um paraffin sections, hybridized with the probe describe above at 40°C for 2 hours. The C3 channel channels was revealed with TSA Opal570 (Akoya Biosciences, Cat. No. FP1488001KT). After 30 minutes blocking with 1% BSA, tissues were incubated with the primary antibody: rabbit anti Olfm4 overnight at 4°C. The secondary antibody: donkey anti rabbit Alexa488 was incubated for 45 minutes at RT before counterstaining with DAPI and coverslipping with Prolong Gold Antifade Mounting medium.

**Aerolysin purification and cell intoxication**
Proaerolysin toxin was produced and purified by our laboratory in *Aeromonas salmonicida* as previously described (*37*). Vero E6 were treated one hour in complete medium with 10ng/ml of proaerolysin at 37°C. Cells were washed twice in complete medium and further incubated at 37°C for indicated times.

**Cell culture methods**
Vero E6 (ATCC: CVCL_0574), HEPG2 (ATCC: HB_8065), Calu-3 (ATCC: HTB_55), HEK (ATCC: CRL_11268) cells were grown in DMEM Media supplemented with 10% fetal calf serum, 1% penicillin-streptomycin. HELA (ATCC: CVCL_0030) cells were grown in MEM Media supplemented with 10% fetal calf serum, 1% penicillin-streptomycin, 1% NEAA and 1% glutamine (all media from ThermoFisher Scientific). HEK293TphACE2-TMPRSS2 cells were kindly provided by Priscilla Turrelly from Didier Trono Lab and cultured as standard HEK293T cells.

**CRISPR/Cas9 deletion**
Vero E6 KO CRIPR-Cas9 for ZDHHC20 were done in the background of Vero E6 WT with the following gRNA sequences to do a deletion in monkey *zDHHC20* gene:
gRNA(F):5’-TGGCGTTAAGCTGATACCATTGG-3’
gRNA(R):5’-CCATTGTGAAGTTAAGACATAGG-3’
Vero E6 KO CRIPR-Cas9 for ZDHHC9 were done in the background of Vero E6 KO ZDHHC20 with the following gRNA sequences to do a deletion in monkey *zDHHC9* gene:
gRNA(R):5’-AGTCTATCCTATCAGCCCACCGG-3’,
gRNA(F):5’-CATTGGGTCACATTTGCGGAAGG-3’

HEK293T KO CRIPR-Cas9 for ZDHHC20 were done in the background of HEK293T WT with the following gRNA sequences to do a deletion in human *zDHHC20* gene:
gRNA (F): 5’-GCGTCCGAGTCACCGTCGCCGGG-3’
gRNA (R): 5’- ATTAAGGCATCATTCTGCTCTGG-3’

**Protein extraction and Western blot**
For Western blot, cells or epithelia or tissues were lysed 30 min at 4°C in IP buffer (0.5% NP40; 500mMtris-HCl, pH=7.4; 20mM EDTA; 2mM benzamidine; 10mM NaF and a cocktail of protease

inhibitors), centrifuge 3 min at 2000 g and protein amount were quantified in supernatants and samples were processed for Western blot with the annotated antibodies.

**Plasmid constructs and overexpression**

Plasmids were transfected in Vero E6 or Hela cells for 24 hours (3ug/9.6cm$^2$ plate using Transit-X2$^{RTM}$ (Mirus). For control transfection, we used an empty pcDNA6.2 plasmid.
Plasmids expressing WT or 10CA SARS-CoV-2 SPIKE were cloned in pcDNA6.2 with HA-tag in C-terminal for initial plasmid (Addgene: 149329). Cystein to alanin substitution were done with Quickchange.
Plasmids expressing ZDHHC20$^{Long}$ and ZDHHC20$^{short}$ were cloned in pcDNA3.1 with myc-tag in N-terminal or with HA-tag in C-terminal. All deletion or mutations to Alanin were done by Quickchange.
Plasmids expressing reporter constructs ZDHHC20$^{Long}$ and ZDHHC20$^{short}$ and mutants were cloned in pcDNA3.1 with all amino acids sequence till the end of first transmembrane domain:**M**WGGGGAWLGAGTGGVPAAPRTFAPTSPASEAGELDQQPPGASESPSPGLPRSPERWD AGLGSRQRN**M**APWTLWRCCQRVVGWVPVLFFITFVVVW**,** followed by a linker**:** RILQSTVPRARDPPVAT and GFP sequence with N-glycolsylation site: MVSKGEELFTGVVPILVELDGDVNGHKFSVSGEGEGDATYGKLTLKFICTTGKLPVPWPTLVTTF TYGVQCFARYPDHMKQHDFFKSAMPEGYVQERTIFFKDDGNYKTRAEVKFEGDTLVNRIELKGI DFKENGSILGHKLEYNYNSHKVYITADKQKNGIKVNFKTRHNIEDGSVQLADHYQQNTPIGDGP VLLPDNHYLSTQSALSKDPNEKRDHMVLLEFVTAAGITLGMDELYKstop. All deletion or mutations to Alanin in reporter constructs were done by Quickchange.
For RUSH experiments, ZDHHC20$^{Long}$ and ZDHHC20$^{short}$ were cloned in RUSH STR-Li-SBP-EGFP plasmids (*24*).
The following plasmids were purchased from Addgene: mScarlet-Giantin_C (85050), FOXA1-flag (153109); SP1-flag (25543).
For stable expression of myc-ZDHHC20 constructs in cells HEK293T-KOZDHHC20, isolated cell clones were selected with 10 µg/ml of puromycin and assessed for equivalent level of expression.

**siRNA and silencing**

Control siRNA, or specific human or Monkey siRNA listed below were purchased from Qiagen and transfected 72 hours with 15pmole/9.6 cm2 plate using Transit-X2$^{RTM}$ (Mirus) as transfection reagent.

| | |
|---|---|
| control: | 5’-ATTGAACAAACGAAACAAGGA-3’ |
| human/monkey ZDHHC9: | 5’-CTCAACCAGACAACCAATGAA-3’ |
| human FOXA1: | 5’-CCAGACGGGTTTCATTATTAT-3’ |
| human FOXA2: | 5’-CACGTTCTATATAAGGAGGAA-3’ |
| human SOX13: | 5’-TTCACAAAGTTTGTTCCCTAA-3’ |
| human RXRA: | 5’-TTCGTGTAAGCAAGTACATAA-3’ |
| human USF: | 5’-CAGAGTAAAGGTGGGATTCTA-3’ |
| human SP1: | 5’-CAGCAAGTTCTGACAGGACTA-3’ |
| human EID1: | 5’-CTCGGCTGTGATGAGATTATT-3’ |
| human GATA1: | 5’-AAGCGCCTGATTGTCAGTAAA-3’ |

**PEGylation- Acyl PEG exchange**

Acyl PEG exchange was used to follow S-acylated protein by addition of mPEG to acylated cysteine following removal of hydroxylamine as previously (*4*). Cell lysates were first incubated 30 min at RT with 10 mM TCEP. Free cysteine in cell lysates were blocked with 100 mM NEM, excess of NEM is removed by acetone precipitation. S-acylated cysteines were revealed by treatment 1 h at 37°C with 200 mM neutral hydroxylamine. Lysates were desalted with Zeba spin culumns and incubated 1 hour at 37°C with 2mM 5kDa methoxypolyethylene glycol maleimide. The addition is stopped by incubation with SDS-PAGE loading buffer with BME, and the lysate is analyzed by SDS PAGE.

**Acyl-RAC**
S-acylated protein were purified using Thiopropyl Sepharose in the presence of 200 mM neutral hydroxylamine from cell lysates pretreated with 100mM NEM to alkylate irrelevant cysteines (*4*).

**Endo H treatment**
Following manufacturer instructions (NEB, P0702S), 40ug of cell extract were denatured 10 minutes at 100°C and treated 1 hour at 37°C with 1000 units of EndoH.

**3H-palmitic acid incorporation-decay**
HeLa cells were transfected with different constructs were incubated 1 hour for starvation in IM (Glasgow minimal essential medium buffered with 10mM Hepes, pH 7.4) and for indicated time in IM with 200 µCi/ml $^3$H palmitic acid (9,10-$^3$H(N)) (American Radiolabeled Chemicals, Inc.). For decay analysis, after starvation, cells were incubated for 3 hours in IM with 200 µCi/ml $^3$H palmitic acid and cells were washed, incubated in DMEM complete medium for the indicated time of chase, or directly lysed for immunoprecipitation with the indicated antibodies.
For immunoprecipitation, cells were washed 3 times PBS, lysed 30 min at 4°C in IP Buffer and centrifuged 3min at 5000 rpm. Supernatants were subjected to preclearing with G sepharose beads prior immunoprecipitation reaction. Supernatants were incubated overnight with the appropriate antibodies and G Sepharose beads. After immunoprecipitation, washes beads were incubated for 5 min at 90°C in reducing sample buffer prior to 4-20% gradient SDS-PAGE. Gels are incubated 30 minutes in a fixative solution (25% isopropanol, 65% H2O, 10% acetic acid), followed by a 30 minutes incubation with signal enhancer Amplify NAMP100 (GE Healthcare). The radiolabeled products were revealed using Typhoon phosphoimager and quantified using the Typhoon Imager (ImageQuanTool, GE Healthcare).

**35S-methionin-cysteine pulse chase**
HeLa cells were starved in DMEM HG devoid of Cys/Met for 30 minutes at 37°C, pulsed with the same medium supplemented with 70 µCi/ml of $^{35}$S Cys/Met (American Radiolabeled Chemicals, Inc.) for 20 minutes, washed and incubated in DMEM complete medium for the indicated time of chase before immunoprecipitation as for 3H-palmitic acid radiolabeling experiments.

**Viral strains, Stock production and titration with plaque-based assays**
Passage 3 (P3) viral stocks were isolated from supernatants of Vero E6 cells, cultured in T75 flasks to a confluency of 80-90%, and infected at MOI≈0.05 with passage-2 (P2) SARS-CoV-2 viral strain (lineage B.1) hCoV-19/Switzerland/GE-SNRCI-29943121/2020, (GISAID ID: EPI_ISL_414019). P2 working Delta or Omicron stocks, were also produced in Vero E6 from Delta B1.617.2 (EPI_ISL_1811202) or Omicron BA.1 (EPI_ISL_7605546) isolates. Infected supernatants containing viral stocks were harvested between 48 and 72 h post-inoculation, approximately 10 ml of DMEM supplemented with 2.5% FCS or Eppiserf serum-free media. Supernatants were clear of cell debris by centrifugation (500 g 10 min) and filtration (0.45 µm), aliquoted, and stored at -80 C. Viral titer was quantified by determining the number of individual plaque-forming units after 48 h of infection in confluent Vero E6 cells. In brief, viral stocks were serially diluted (10-fold) in serum-free medium and (400 µl) inoculated in triplicate 48 wells, confluent Vero E6 cells (2.5 x 105) cells per well. After 1 h, inoculums were discarded, and cells were overlaid with a mixture of 0.4% of Avicel-3515 (Dupont) (from 2% stock) in DMEM supplemented with 5% FCS and penicillin and streptomycin for an additional 48 h. Overlays were discarded, and cells were fixed in 4% paraformaldehyde (PFA) for 30 min at RT. Fixed cells were washed in PBS and stained with 0.1% crystal violet solution (in 20% ethanol/water) for 15 min. The staining solution was discarded, and the wells were washed twice in water. Plates were allowed to dry and analyzed for quantification of the cytopathic effect as the number of individual Plaque forming units (PFU) per ml (Avg PFU*1/Volume*1/dilution factor). Such un-concentrated viral stocks yielded between 0.5 to 5x106 PFU/ml.

**SARS CoV-2 infections**
Unless otherwise indicated, all infections were done using P3 SARS-CoV-2 (B.1) stocks. P2 stocks were used for experiments using the Omicron and Delta variants. Vero E6, Calu-3 cells or HEK293T seeded to a confluency of 90 to 100% were, washed twice in warm serum-free medium and inoculated with the indicated MOI of SARS-CoV-2, diluted in serum-free medium (5 ml for a T75 and 2 ml for T25 flask – stocks and biochemistry experiments, and 500 µl for 12-well plates - infectivity assays). 1 hour after inoculation, cells were washed with complete medium, and infection was allowed to proceed for the indicated time points in DMEM supplemented with 2.5% FCS, penicillin, and streptomycin (10 ml for T75; 4 ml for T25, and 1 ml for 12 well plates). Infected cells and virion supernatants were harvested and inactivated in IP buffer (1:1 v/v for supernatants) or lysed in Pegylation/Acyl Rac lysis buffer (0.5% Triton-X100, 2.5% SDS, 25 mM HEPES, 25 mM NaCl, 1 mM EDTA, pH 7.4, and protease inhibitor cocktail) for at least 30 min.

**Titration of viral E and N copies**
For titration of viral RNA, equivalent volumes of RNA extracted from infected culture supernatants or RNA from serial dilutions of SARS-CoV-2 (N+E) RNA Quant Standard (Cat.# AM2050 Promega) were used for cDNA synthesis. Samples were used to perform QPCR analysis (using E and N specific primers) as described. A standard curve was generated by plotting the Ct values against the number of E/N copies per µl indicated by the serial dilution of the E/N standards (stock at 4 × 106 copies/µl) and used to extrapolate the number of Viral N/E copies/ml in samples.

**SARS-CoV-2 single round infectivity analysis**
KO-ZDHHC20/9 cells cultured in T75 flasks transfected with empty plasmids or ZDHHC20 (Short or Long) expressing plasmids (24 h) were infected as described above. Supernatants were harvested between 24 and 48 h post inoculation and processed for QPCR analysis (150 µl) or aliquoted and stored at -80°C until titration of viral N/E copies. After titration and adjustment to viral N/E RNA copies non-concentrated SARS-CoV-2 supernatants were used to infect confluent Vero E6 cell monolayers cultured in 12 well plates using an approximate ratio of 5 to 10 N/E copies per cultured host cell. At least 3 wells were infected per supernatant per condition. Infection was done as described until 6 h post inoculation when cells were washed lysed and processed for QPCR analysis. Infected cells were harvested and lysed in 330 µl of Maxwell® RSC Viral Total Nucleic Acid Purification Kit-lysis buffer from Promega, incubated at 80°C for 10 min, and used for Viral RNA extraction according to manufacturer's instructions. RNA concentration was measured, and 500 ng or 1000 ng of total RNA was used for cDNA synthesis using iScript. A 1:5 dilution of cDNA was used to perform quantitative real-time PCR (QPCR) as described below. Ct values from primers used for viral genes (E, RdRp, and N) were used for quantification of N/E copies using SARS-CoV-2 RNA standards as described or normalized to host housekeeping genes (ALAS-1, RPL27) for cell replication infectivity assays. Results were expressed as $2^{(-\Delta\Delta Ct)}$*100%.

**VLPs production**
HiBiT-N-tagged virus-like particles (VLPs) were produced as described previously (*4*, *38*). Briefly, equimolar amounts of full-length CoV S, E (envelope), M (membrane), and HiBiT-N-encoding plasmids (total, 10 µg) were transfected into KO-ZDHHC20 HEK293T cells stably expressing plasmids coding for ZDHHC20 (short or Long) or empty plasmids) and co-transfected with siRNA targeting ZDHHC9 for 72 h. To produce Bald ("no-S") VLPs, the S expression plasmids were replaced with empty vector plasmids. At 6 h post-transfection, cells were replenished with fresh DMEM-10% FBS. HiBiT-N VLP suspensions were collected in FBS-free DMEM or Eppiserf from 24 to 48 h post-transfection. Suspensions containing HiBiT-N VLPs were clarified by centrifugation (300 g, 4°C, 10 min; 3,000 g, 4°C, 10 min). To obtain purified viral particles, clarified VLP suspensions were concentrated 100-fold by overlaid onto 20%, wt/wt, sucrose cushions and particles purified via slow-speed pelleting (SW32 8000 rpm, 4°C, 20 h). The resulting pellet was resuspended in Eppiserf to 1/100 of the original medium volumes. VLPs were stored

at -80°C or analyzed promptly for titration using Nano-Glo® HiBiT Extracellular Detection System (Promega #N2420) with passive lysis buffer (Promega #E1941).

**VLP cell entry assay**

HEK293TphACE2-TMPRSS2 target cells, cultured in 96 well plates precoated with poly-lysine, were transfected with pcDNA3.1-hACE2-LgBit. At 2 days post-transfection, cells were incubated with a live-cell Nluc substrate (Nano-Glo Vivazine; Promega), and 2 h later, cells were incubated at 4°C on ice. HiBiT-N VLPs were inoculated at equivalent N-HiBiT input multiplicities, using 4 independent wells per VLP stock replicate,. Nluc levels were quantified immediately and set as T=0, before incubation of the plates at 37°C. HiBiT-N VLPs lacking Spike proteins (bald) served as negative controls. At the indicated intervals following VLP inoculation, Nluc levels were quantified using a HIDEX microplate reader. For data presentation, the Nluc recordings in cultures inoculated with bald VLPs were normalized to values of 1.0, and the fold increases over this control condition were calculated and plotted as "relative entry."

**Human MucilAir viral assays**

MucilAir-human upper respiratory tissues (Epithelix, Geneve, Switzerland) were maintained according to the manufacturer's protocol. Prior infection, tissues were washed apically with 200 µl of DPBS, calcium, magnesium (#14040091 – Gibco) for 20 min at 37°C and the basal medium replaced with fresh mucilair medium. Tissues were infected at the MOI of 0.1 PFU (assuming the manufacture's estimations of 500000 cells per tissue). Tissues were inoculated with 200 µl of SARS-CoV-2 (passage 3) for 3 h (apically) at 33°C. Apical inoculum was removed and infected tissues maintained for until harvest. Harvest was performed using 300 µl lysis buffer for 30 min. Lysed inactivated tissue lysates were transferred to clean tubes and processed for Western blot.

**hK18- ace2 mice-Virus infection**

Eleven-week-old male and female K18-hAce2 C57BL/6j transgenic mice (strain: 2B6.Cg-Tg(K18-ACE2)2Prlmn/J) from The Jackson Laboratory were administrated intranasally $1x10^4$ PFU SARS-CoV-2 (B.1). Mice were euthanized after 5 or 6 days of infection and immediately dissected for organs collection and further analysis. Proteins were extracted as described above and viral and tissue RNA were extracted following instruction of Omega Bio-TEK, (EZNA total RNA KIT I). Procedures were performed according to protocols approved by the Veterinary Authorities of the Canton Vaud and according to the Swiss Law (license VD3794A, EPFL).

**DSS induced mice colitis**

Eight-weeks CMG2+/+ or CMG2 -/- male were given 3% Dextran Sulfate Sodium in the drinking water for 7 days, then switched to regular drinking water for 3 days. Control Eight-weeks CMG2+/+ or CMG2 -/- male were given drinking water that did not contain DSS. On day 10, mice were euthanized with injection of pentobarbital and bled via cardiac puncture and colon and intestines were collected. For animal experimentation, all procedures were performed according to protocols approved by the Veterinary Authorities of the Canton Vaud and according to the Swiss Law (license VD 3497, EPFL).

**Quantitative real-time PCR**

RNA was extracted from cells using Maxwell RSC Promega Kit (AS1330) and from mice tissues using E.Z.N.A total RNA kit (R6834). RNA concentration was measured and 500 ng of total RNA was used for cDNA synthesis using iScript (Biorad : 1708891). A 1:5 dilution of cDNA was used to performed quantitative real-time PCR using Applied Biosystems SYBR Green Master Mix (Thermofisher Scientific) on 7900HT Fast QPCR Applied Biosystems with SDS 2.4 Software.

The data in triplicate were normalized to housekeeping genes (human: ALAS1, GUSS, TBP; mouse: RSP9, EEF1A1,COX6a1). Results were expressed as 2^(-DDCt)*100%.

Primers (all 5' to 3') used were:

Housekeeping gene:
Human GUSS : F: CCACCAGGGACCATCCAAT ; R: AGTCAAAATATGTGTTCTGGACAAAGTAA
Human TBP: F: GCCCGAAACGCCGAATATA ; R : CGTGGCTCTCTTATCCTCATGA
Human ALAS-1: F : CTCACCACACACCCCAGATG; R : AGTTCCAGCCCCACTTGCT
Human RPL27: TGTCCTGGCTGGACGCTACT; CTGAGGTGCCATCATCAATGTT
Mouse RSP9: F: GACCAGGAGCTAAAGTTGATTGGA; R: TCTTGGCCAGGGTAAACTTGA

Mouse EEF1A1: F: TCCACTTGGTCGCTTTGCT; R: CTTCTTGTCCACAGCTTTGATGA

Mouse COX6a1: F: CTCTTCCACAACCCTCATGTGA ; R : GAGGCCAGGTTCTCTTTACTCATC
Human ZDHHC20:
Human C1: F: CGCCTTGTGGGGATGGATCC; R: CCGCTTGTTGGACAGTGAATCTCAG
Human C2: F: GTCGTCTGGTCCTACTACGC; R: AGCCACAAGGTAAACAACGG
Human 1: F: GTGGCCTTATTTGGAAGTGGG; R: CCCTACTCTAGTATGGCCTC
Human 2: F: GCCACTGCCACTGCAGGTC; R : CACAGCCATGTGCCCTCTG
Human 3: F: GGCACAAAATCAGGGAGAACAG; R: AAGAAGACGTCACCCTTTGCT
Human 4: F: AGTCTCCCTCCCCTATTGAGT; R: AAAACACGCCCTTGATGGAT
Human 5: F: CCTCGGACTTTTGCTCCCACAAG; R: CGTCCCACCGTTCTGGGGAG
Mouse ZDHHC20:
Mouse C1: F: ATATTGCCTTTTTGTGGCTGC; R: ACTGTTGGTTCATTCGTCCAA
Mouse C2: F: TCATCACTGTCCATGGGTGAA; R: ACTGTTGGTTCATTCGTCCAA
Mouse 5'UTR : F: CGCCCTTCGCCACTGCTTG; R: CTGGGGTCTTGTGGTCCTAC
Human and Monkey Transcription Factors
FOXA1: F: TGGAACAGCTACTACGCAGAC; R:GGTGTTCATGGTCATGTAGGTG
SP1: F:GCCACCATGAGCGACCA; R: GAAAAGGCACCACCACCATT
EID1: F: TCGTCTGACCGAAGAACTCG; R: TGGGTCCCTCCTCAAGTAGT
SOX13: F: CTCCAGAGGGTAATGGGTCC; R: CTATGGCTGGCACCACTTCT
USF1: F: CCTTGGATAGGAAAGGACTTAGC; R: ATCTGCACTGTCCCCTCTTC
RXRA: F: CAGCGGAACCAAAACTGCT; R: GGTGAGCTGAGCCGGT
Mouse Transcription Factors
FOXA1: F: ACTCTCCTTATGGCGCTACC; R: ACACCTTGGTAGTAGGCTGG
SP1: F: GTGGGAAGCGCTTTACACGTTCGG; R: GCCTGCCCTGAGTGCCCTAAG
SOX13 : F: CCCGACCGATTAGATGTCCA; R: GCAAGGCTCCTTCTTCTCCT
USF1 : F: ATCCAAAGACGGAGAAGGCT; R: GAATGCTAAGTCCGGGCCA
RXRA : F: GCAGACATGGACACCAAACA; R: CACCTGGGTAGAGAAGTCGAG

SARS-CoV-2
E SARBECO: F: ACAGGTACGTTAATAGTTAATAGCGT; R: ATATTGCAGCAGTACGCACACA
RDRP: F: AGCTTGTCACACCGTTTC ; R: AAGCAGTTGTGGCATCTC
N (Nucleocapsid): F: GACCCCAAAATCAGCGAAAT;R: TGTAGCACG ATTGCAGCATTG

**5'-rapid amplification of the cDNA ends (5'-RACE)**

We applied SMARTer 5'-RACE technique using independent RNA preparations extracted from Calu-3 cells infected with SARS-CoV-2 for 24 h at MOI 0.1. A modification of the manufacturer's protocol (Clontech) was carried in order to generate first-Strand cDNA Synthesis from specifically *zdhhc20* transcripts. For that, instead of using the modified oligo (dT) primer, cDNA synthesis was done using a *zdhhc20* specific reverse primer: R:GTA CGC GTA GTA GGA CCA GAC. 5'RACE was then carried with the gene (*zdhhc20*) specific primer #4 modified for infusion cloning. 5' GAT TAC GCC AAG CTT CGT CCC ACC GTT CTG GGG AG - 3'. 5'-RACE products were loaded in an agarose gel and a single large fragment around >9000 bp was excised for DNA purification and In-Fusion cloning (Clontech). A total of 53 transformed clones were confirmed as *zdhhc20* 5'ends by Sanger sequencing using *zdhhc20* primer #4. A summary of the sequenced mRNA species and their frequency is shown in fig S1D and Table S1.

**Immunohistochemistry**

Vero E6 or HeLa cells seeded in glass coverslips in 24-well plates and transfected with the indicated myc-/3HA-/GFP-ZDHHC20 constructs (for 24 h) were fixed in 4% paraformaldehyde (15 min), quenched with 50 mM NH4Cl (30 min) permeabilized or not with 0.05% Saponin (5 min). Antibodies were diluted in PBS containing 1% BSA and 0.05% Saponin for permeabilization. Coverslips blocked in PBS 1% BSA, for 30 min, washed and incubated with primary antibodies overnight at 4°C, washed three times in PBS, and incubated 45 min to 1 h with secondary antibodies and when indicated nuclear staining Hoechst. Coverslips were mounted onto microscope slides with ProLong™ Gold Antifade Mountant. For exclusive imaging of GFP-expressing reporters, coverslips were nuclear-stained and mounted without permeabilization. Images were collected using a confocal laser-scanning microscope (Zeiss LSM 700) and processed using Fiji™ software.

**Rush**

Cells were seeded in 35-mm glass bottom plates (FD35-PDL, FluoroDish™) and transfected with 1 µg of each plasmid DNA encoding for ZDHHC20-RUSH reporter and Scarlet-Giantin. RUSH reporters encode for both a GFP-ZDHHC20 fusion tagged with streptavidin binding protein (luminal) and the ER resident hook - the transmembrane invariant chain (Ii) – streptavidin tagged. After 48 h of transfection, cells were washed and culture medium replaced with pre-warmed carbonate independent Leibovitz's medium (Invitrogen). Time-lapse imaging was performed on a Visitron Spinning Disk CSU W1 with full temperature control and CO2, under culture conditions. Z-stack (0.7 µm slice) images were acquired for each channel before and after addition of Bio-D (T=0) at final concentration 40 µM, every 10 s for at least 1800 s (30 min). Acquisitions were processed using Fiji™ software 45. The mean fluorescence intensities were measured for each time-lapse image, for two ROIs: Total cell (marked by initial GFP-ZDHHC20 staining) and Golgi (marked by Scarlet-Giantin). Golgi/total fluorescent ratios were normalized to 1 for T=0 and corrected values plotted using GraphPad Prism.

**Automated microscopy analysis**

Approximately 10000 Hela MZ cells were plated in FluorobriteTM in 96 wells imaging plates from Ibidi (ref: 89626) for 24 h. Cells were transfected for 24 h with 1 µg/mL final concentration of different PAT20 plasmids using Mirus Transit-X2 transfection reagent at a final dilution of 1/1000. Cells were fixed with 3% PFA, permeabilized with 0.05% saponin and blocked with 1% BSA. Cells were then stained with anti GM130 and anti MYC (when necessary) antibodies both at 1/200 followed by incubation with secondary antibodies tagged with alexa-fluor 647 and 488 (for MYC staining) with Hoechst at 1/2500. In all conditions at least 9 images per well were acquired using IXM confocal automatic microscope (Molecular Device) using 20X water immersion objective 0.95NA. Analysis of the automated microscopy images was done via the MetaXpress Custom Module editor software. The images were segmented to generate relevant masks, which were then applied on the fluorescence images to extract the relevant measurements. Cell and Nuclei masks were created using Hoechst to generate the master object (cell). To facilitate segmentation of Golgi, we applied the top hat deconvolution method, to reduce the background noise and highlight bright objects. A logical operation was used to generate the mask of cytoplasm without Golgi. Relevant masks were then applied on the fluorescent images to extract relevant measurements.

**Phyml tree**

A list of orthologs from all species for the human gene ZDHHC20 using the ensembl.org web site was established. These transcripts were used as query sequences to a blastn/megablast search to retrieve RefSeq and EMBL/Genbank transcripts with very high similarity.
Those retrieved transcripts were analyzed using a Perl script to find potential 5' extension of the known ORF until the 5'most potential TSS while staying in-frame for the transcription. All the retrieved transcripts (both those where an extension is possible and those where it is not) were translated to the corresponding amino acid letter codes and aligned using mafft into a multiple sequence alignment (MSA). The MSA was hand-edited to remove duplicates and select one representative sequence per broad taxonomic group. The taxonomic information was retrieved from the corresponding RefSeq and/or EMBL/Genbank entries.
The resulting MSA was provided as input to phyml, which was requested to perform 100 cycles of bootstrap. The resulting output tree was plotted using the ape R package to produce a PDF document that was used to generate the figure.

**Reproducibility and statistical analysis**

Unless otherwise indicated, all data were repeated at least three times independently. Unless otherwise stated, each data point corresponds to one independent experiment with consistent results. Statistical analysis was carried out using Prism software. Data representations and statistical details can be found in the description of the figures. For ANOVA analysis, p values were obtained by post hoc tests to compare every mean and pair of means (Tukey’s & Sidak’s) or to compare every mean to a control sample (Dunnet’s). Estimated half-lives, from metabolically labelling experiments were extracted from the individual experiments using non-linear regression with one-phase decay. (Prism).

**Fig. S1.**

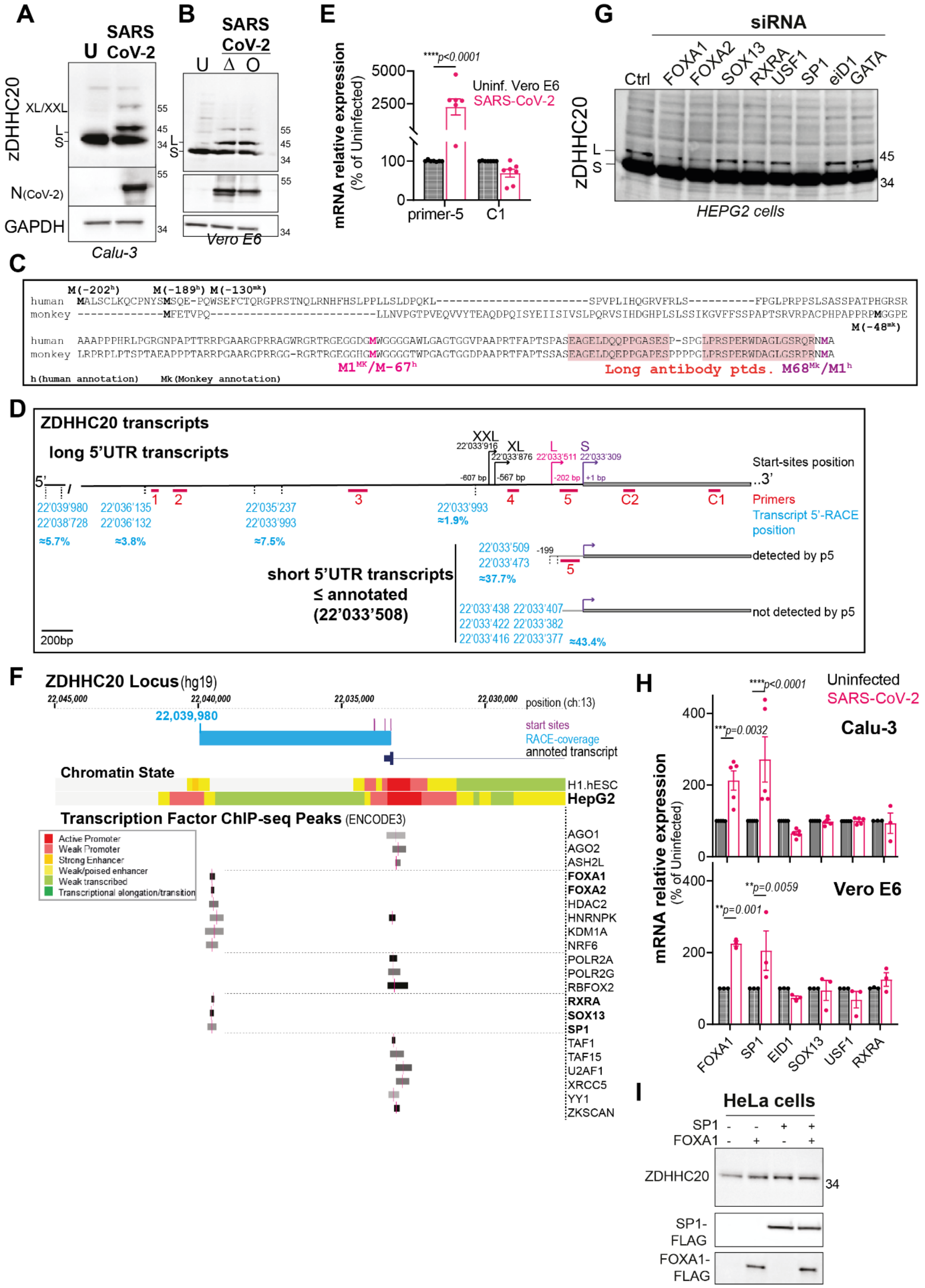

A
SARS CoV-2
U
zDHHC20
XL/XXL
L
S
55
45
34
N(CoV-2)
GAPDH
Calu-3
B
SARS CoV-2
U Δ O
Vero E6
E
mRNA relative expression (% of Uninfected)
****p<0.0001
Uninf. Vero E6
SARS-CoV-2
5000
2500
100
0
primer-5
C1
G
siRNA
Ctrl
FOXA1
FOXA2
SOX13
RXRA
USF1
SP1
eID1
GATA
HEPG2 cells
C
M(-202h) M(-189h) M(-130mk)
M(-48mk)
M1MK/M-67h
Long antibody ptds.
M68Mk/M1h
h(human annotation)
Mk(Monkey annotation)
D
ZDHHC20 transcripts
long 5'UTR transcripts
XXL
XL
L
S
22'033'916
22'033'876
22'033'511
22'033'309
-607 bp
-567 bp
-202 bp
+1 bp
Start-sites position
Primers
Transcript 5'-RACE position
22'039'980
22'038'728
22'036'135
22'036'132
22'035'237
22'033'993
≈5.7%
≈3.8%
≈7.5%
≈1.9%
1
2
3
4
5
C2
C1
short 5'UTR transcripts ≤ annotated (22'033'508)
22'033'509
22'033'473
-199
≈37.7%
detected by p5
22'033'438 22'033'407
22'033'422 22'033'382
22'033'416 22'033'377
≈43.4%
not detected by p5
200bp
F
ZDHHC20 Locus (hg19)
22,045,000
22,040,000
22,035,000
22,030,000
position (ch:13)
22,039,980
start sites
RACE-coverage
annoted transcript
Chromatin State
H1.hESC
HepG2
Transcription Factor ChIP-seq Peaks (ENCODE3)
Active Promoter
Weak Promoter
Strong Enhancer
Weak/poised enhancer
Weak transcribed
Transcriptional elongation/transition
AGO1
AGO2
ASH2L
FOXA1
FOXA2
HDAC2
HNRNPK
KDM1A
NRF6
POLR2A
POLR2G
RBFOX2
RXRA
SOX13
SP1
TAF1
TAF15
U2AF1
XRCC5
YY1
ZKSCAN
H
mRNA relative expression (% of Uninfected)
Uninfected
SARS-CoV-2
Calu-3
****p<0.0001
***p=0.0032
Vero E6
**p=0.0059
**p=0.001
400
200
0
FOXA1
SP1
EID1
SOX13
USF1
RXRA
I
HeLa cells
SP1
FOXA1
ZDHHC20
SP1-FLAG
FOXA1-FLAG
34

**Figure S1**

- **A.** WB of ZDHHC20 $^{ALL}$, N, GAPDH on Calu-3 cell extracts uninfected (U) or infected 24h with SARS-CoV-2, MOI=0.1.
- **B.** WB as in A of HEK293T-TMPRSS2-ACE2 extracts uninfected (U) or infected 24 h with SARS-CoV-2 strain B.1 (left), or SARS-CoV-2 Delta B1.617.2 or Omicron BA.1 (right).
- **C.** Alignment of N-terminal amino acids of ZDHHC20 in human (Q5WOZ9), and Green Monkey Chlorocebus Sabaeus (A0A0D9RZN5). First Methionin M1 ($M1^{H}/M68^{MK}$) annotated in human in Magenta. First Methionin annotated in green monkey in pink M1 ($M1^{MK}/M\text{-}67^{H}$), others methionine in-frame in 5' in black. Peptides used to generate rabbit antibodies against Long ZDHHC20 are highlighted in orange.
- **D.** Schematic representation of the 5'UTR from *zdhhc20* transcripts. Long, transcripts > than annotated; Short ≤ annotated 5'UTR. 5' ends obtained by 5'RACE analysis and corresponding abundance indicated in blue (see table S1). Position of QPCR primers in *zdhhc20* locus used in Fig 1G (red) and different transcription start sites are indicated.
- **E.** mRNA quantification using primers probing for different locations in z*dhhc20* transcripts (coding region: C1 or in 5'UTR: primer-5) in Vero E6 cells treated as in A.
- **F.** Overview of UCSC genome browser at *zdhhc20* locus and zoom in at its transcription start site (TSS) (Human version hg19). Two custom tracks and three UCSC genome browser tracks are shown for this region: 1) start-sites (ATGs) indicates *zdhhc20* in-frame ATG codons; 2) 5'-RACE Sequence describes the genomic region amplified by 5'-RACE technique; 3) Transcript annotation corresponds to GENCODE Genes track (version V4 0lift37); 4) Chromatin State displays chromatin state segmentation data from ENCODE consortia for two different cell lines (H1.hESC and HepG2) 5) Transcription Factor (TF) CHIP-seq Peaks track shows transcription factor (TF) binding sites for HepG2 cell line based on ChIP-seq experiments from ENCODE. Only TF with a minimum score range of 300 are shown. The level of peak enrichment is represented with the darkness of the item, and the vertical pink bar marks the point-source of the peak. TFs that have been experimentally tested in this study are shown in pink.
- **G.** WB of ZDHHC20 on HEPG2 cell extracts transfected 72 hours with siRNA targeting the indicated TFs
- **H.** mRNA quantification of the indicated TFs in Calu-3 cells or Vero E6 cells processed as in A.
- **I.** WBs of ZDHHC20 and Flag on Hela cell extracts transfected 24 hours with plasmids expressing SP1-Flag or FOXA1-Flag.

**Fig. S2.**

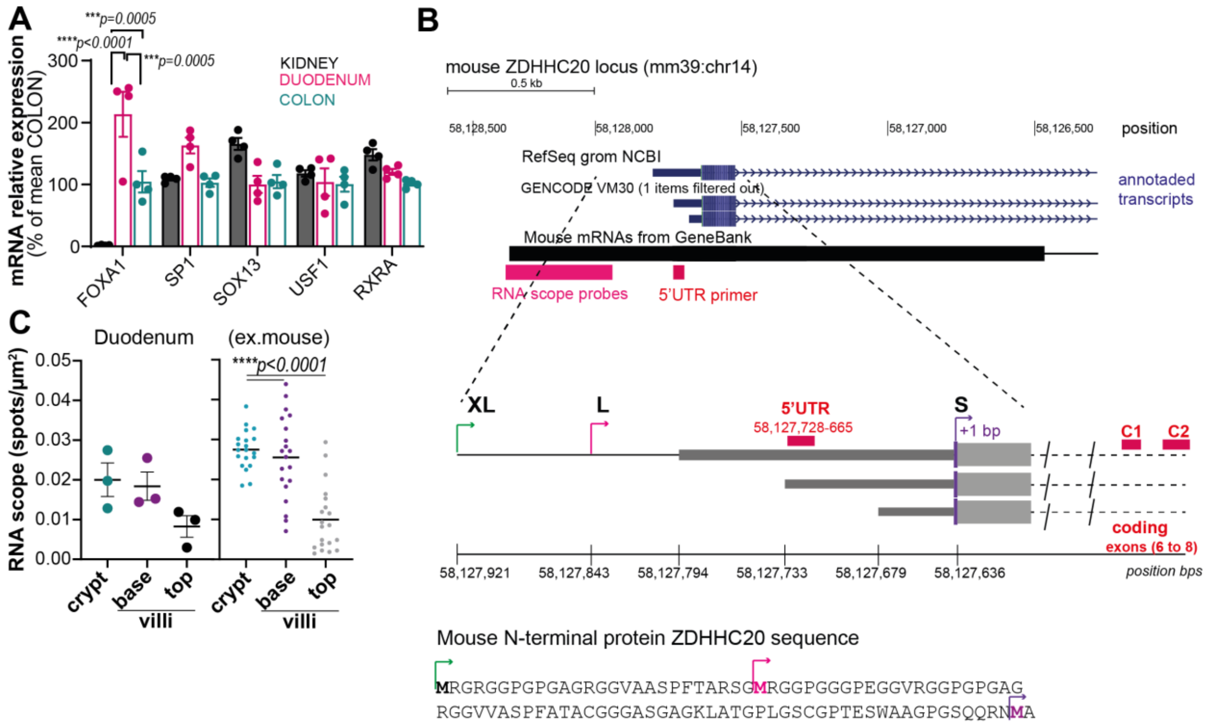


### Supplementary Figure S2

- **A.** mRNA quantification in mouse kidney, duodenum and colon tissues, using primers for the indicated TFs. Results are mean + SEM and each dot represents one of 4 independent mice. *P* values were obtained by two-way ANOVA with Tukey's multiple comparison.
- **B.** Overview of murine *zdhhc20* locus (Mouse GRCm39/mm39) from UCSC genome browser https://genome.ucsc.edu) showing tracks corresponding to transcripts from NCBI RNA reference sequences-Refseq and GENCODE (version VM30) (blue) and coverage of publicly available mouse *zdhhc20* mRNA sequences from Genebank (black). The position of *zdhhc20* RNA scope probes used in Fig 2B (pink) and QPCR primers (red) used in Fig 2E are indicated. Insets show the position of in-frame ATG codons at transcription start sites coding for XL (green), Long (magenta) and Short ZDHHC20 forms with the correspondent N-terminal extension depicted below.
- **C.** Average (left) and representative (one mouse) quantification of *zDHHC20* RNAscope spots on different parts (crypt, base, top) of the mouse duodenum**.** Average results (left) are mean + SEM and each dot represents one of 3 independent mice, whereas representative data (right) depicts mean values and corresponding distribution of RNA scope spots per randomly selected field-of-views (n = 20) for one representative mouse. *P* values were obtained by One-way ANOVA with Tukey's multiple comparison.

**Fig. S3.**

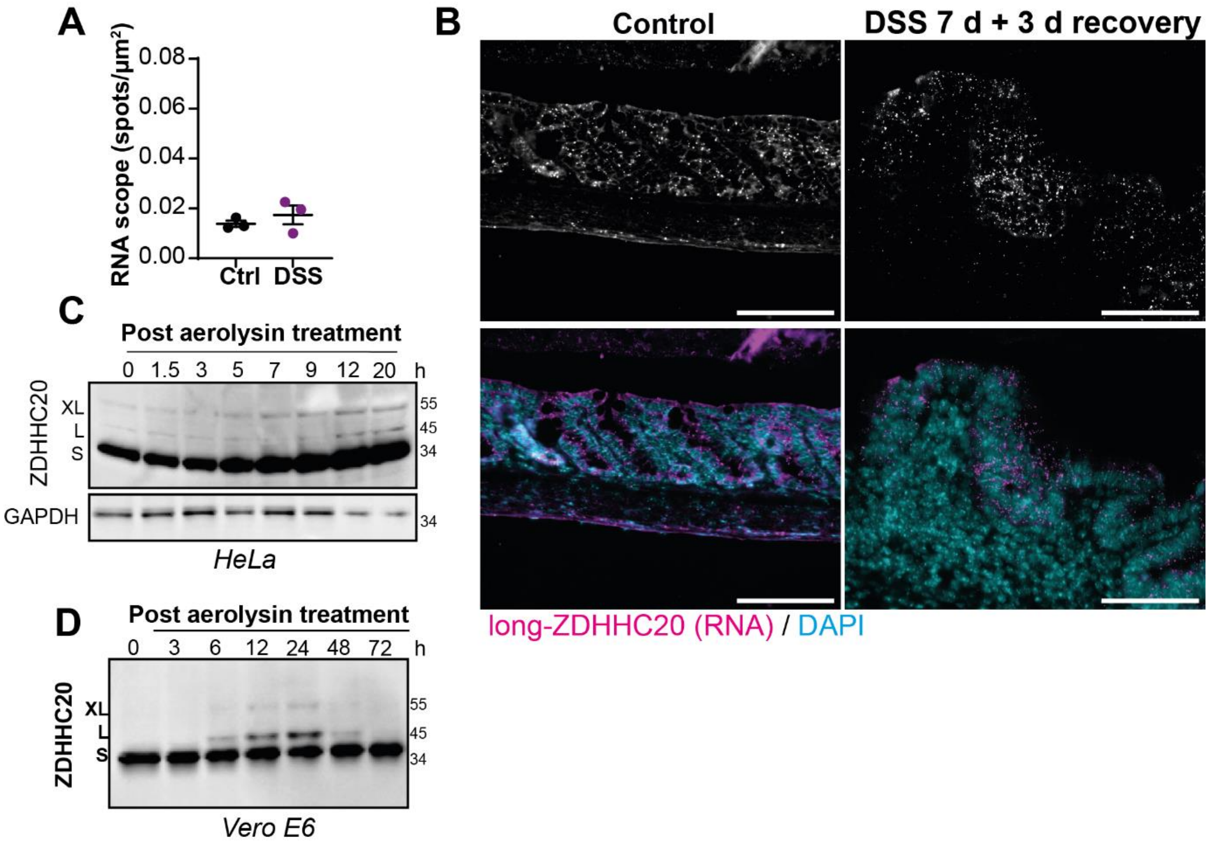


**Supplementary Figure S3**

- **A.** Quantification of ZDHHC20 RNA scope spots on mouse colon from control or DSS-treated mice (7 days treatment plus 3 days recovery). Results are mean + SEM of n = 3 independent mice per condition
- **B.** Representative mouse colon sections of mice treated as in A labelled with RNAscope probe for 5'UTR *zDHHC20* RNA, and stained with DAPI. Scale bars 100 µm.
- **C.** **D.** WB of ZDHHC20 expression from, **C-**HeLa or, **D**-Vero E6 cell extracts treated 1 h with 10 ng/ml of proaerolysin at 37°C, washed and further incubated at 37°C for indicated time-points. ZDHHC20 S, L and XL forms are indicated and GAPDH was used as loading control in C.

**Fig. S4.**

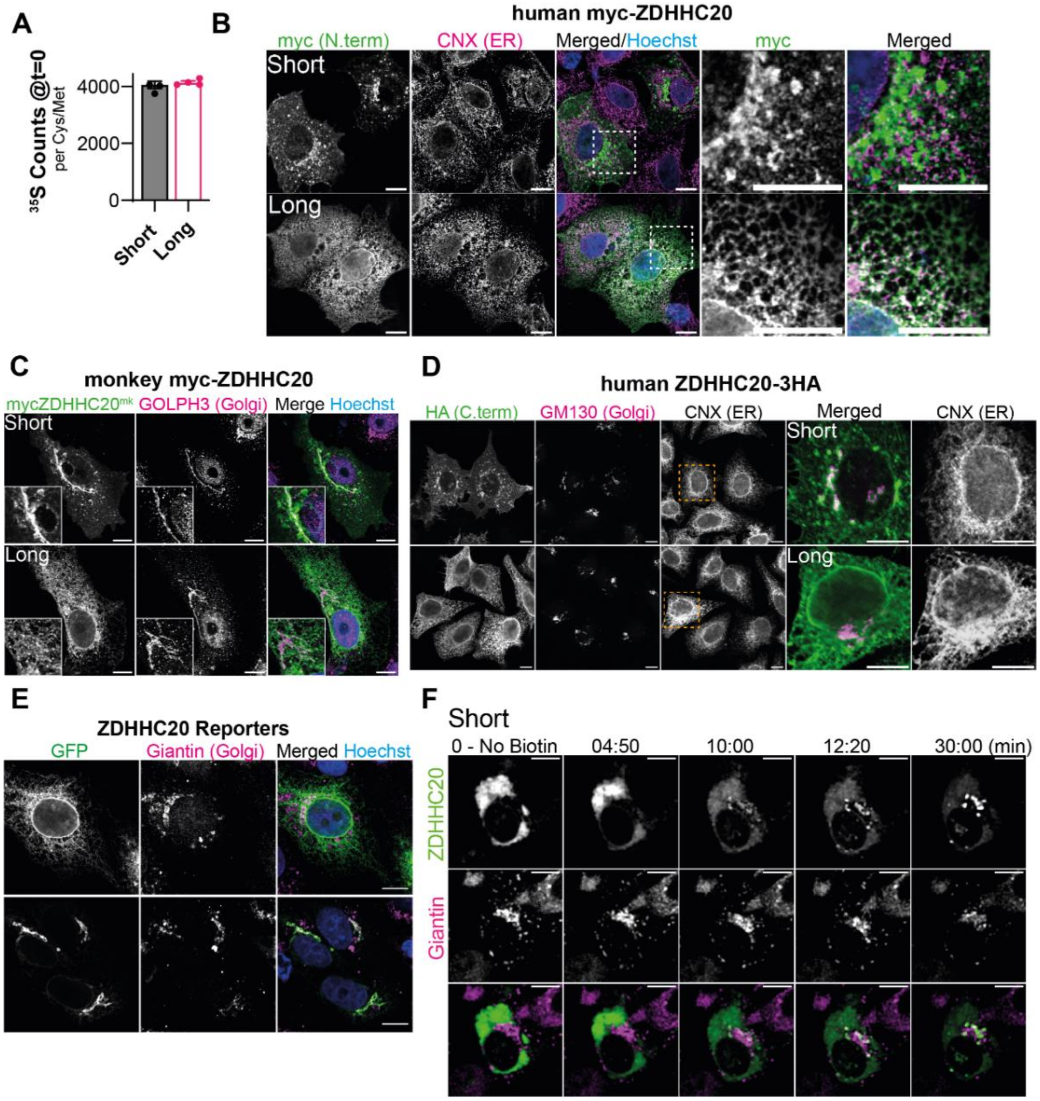
A
B
$^{35}$S Counts @t=0
per Cys/Met
4000
2000
0
Short
Long
human myc-ZDHHC20
myc (N.term)
CNX (ER)
Merged/Hoechst
myc
Merged
Short
Long
C
monkey myc-ZDHHC20
mycZDHHC20mk
GOLPH3 (Golgi)
Merge
Hoechst
Short
Long
D
human ZDHHC20-3HA
HA (C.term)
GM130 (Golgi)
CNX (ER)
Merged
CNX (ER)
Short
Long
E
ZDHHC20 Reporters
GFP
Giantin (Golgi)
Merged
Hoechst
F
Short
0 - No Biotin
04:50
10:00
12:20
30:00 (min)
ZDHHC20
Giantin

**Supplementary Figure S4**

**A.** Hela cells expressing myc-ZDHHC20 (Short or Long) metabolically labelled with 35S-Met/Cys for 20 min. 35S incorporation in myc-ZDHHC20 immunoprecipitation fractions were normalized by the number of Cys/Met residues per ZDHHC20 (20 for short and 21 for Long) and are mean ± SD of n = 4 independent experiments.

**B.** IF of Vero E6 cells expressing myc-ZDHHC20 $^{\mathbf{Short}}$ or myc-ZDHHC20$^{\mathbf{Long}}$ labelled for myc, the ER marker Calnexin (CNX), and nuclear-stained with Hoechst. Scale bar: 10µm.

**C.** IF of Vero E6 cells expressing myc-ZDHHC20 $^{\mathbf{Short}}$ or myc-ZDHHC20$^{Long}$ (Green Monkey Chlorocebus Sabaeus-A0A0D9RZN5 sequence) labelled for myc the Golgi marker (GOLPH3), scale bar: 10 µm.

**D.** IF of HeLa cells expressing C-terminally tagged ZDHHC20$^{\mathbf{Short}}$ -3HA or ZDHHC20$^{Long}$ -3HA labelled for HA, Golgi marker GM130 and ER marker CNX, scale bar: 10µm.

**E.** IF of Vero E6 cells expressing ZDHHC20$^{Short}$- or ZDHHC20$^{Long}$-reporters labelled for Golgi marker Giantin and nuclear-stained with Hoechst, scale bar: 10 µm.

**F.** Time-lapse confocal microscopy images of Vero E6 cells co-expressing (24 h) the Golgi marker Scarlet-Giantin, ZDHHC20$^{short}$-RUSH-GFP reporter and an ER-resident hook. Synchronized trafficking was monitored upon D-biotin addition (after $T_0$ see methods, Fig 4L and movies S1-2).

**Fig. S5.**

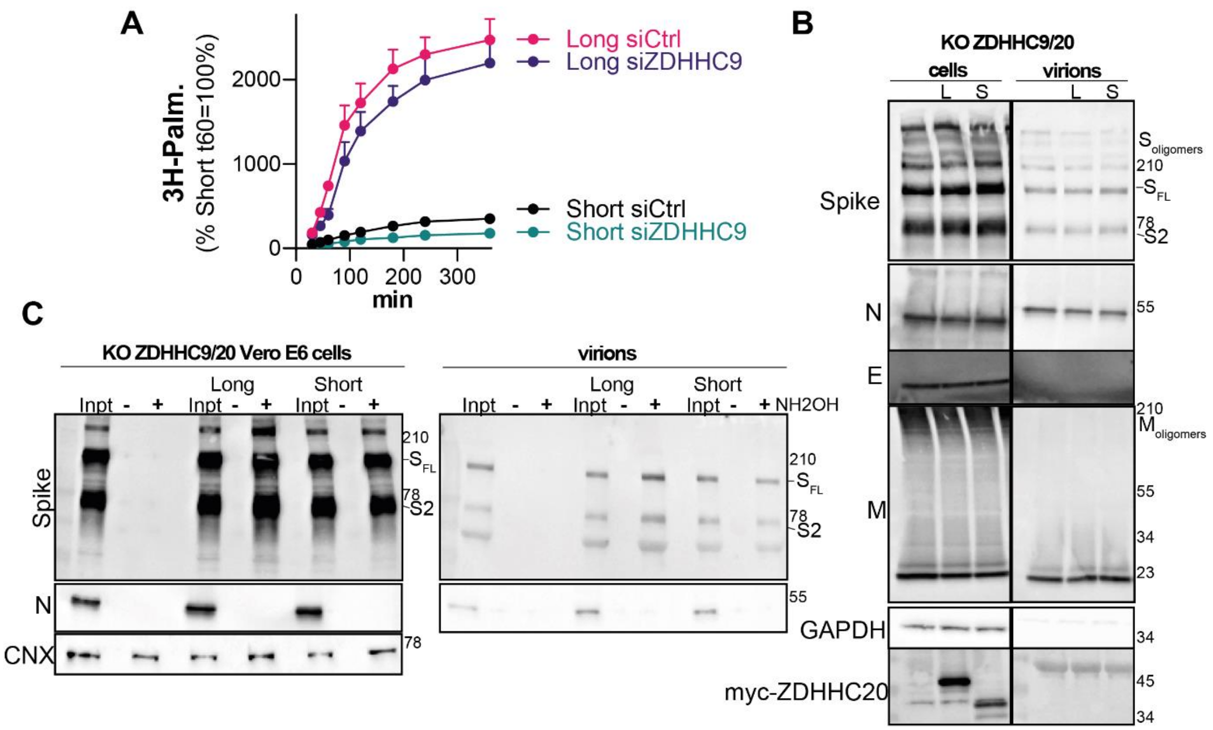


**Supplementary Figure S5**

A. Spike-HA incorporated 3H-palmitic acid normalized to T=60min in Vero E6 KO for ZDHHC20 with or without 3 days siRNA for ZDHHC9 recomplemented with ZDHHC20 $^{\mathbf{Short}}$ or ZDHHC20 $^{\mathbf{Long}}$ with half amount of DNA (eq.protein).
B. Western Blots of Spike, N, E, M, GAPDH, MYC-ZDHHC20 on Vero E6 KO ZDHHC9/20 cell extracts (cells) transfected 24h with control (Ctrl), ZDHHC20$^{Short}$ (Short) or ZDHHC20$^{Long}$ (Long) expressing plasmids, infected 24h with SARS-CoV-2, MOI:0.1, or on virions extracts collected from the corresponding cells.
C. Acyl-RAC on Vero E6 KO ZDHHC9/20 cells described in S5B, blotted for Spike, N and positive control Calnexin (CNX).

**Table S1.**

| 5 ' end positions determined by 5'RACE | Number of clones retrieved | position in relation to start site (-bp) | primer coverage | % | Sum % in fig S1D |
|---|---|---|---|---|---|
| 22,039,980 | 2 | 6671 | all | 3.8 | |
| 22,038,728 | 1 | 5419 | all | 1.9 | 5.7 |
| 22,036,135 | 1 | 2826 | all | 1.9 | |
| 22,036,132 | 1 | 2823 | all | 1.9 | 3.8 |
| 22,035,430 | 2 | 2121 | 3,4,5 | 3.8 | |
| 22,035,237 | 2 | 1928 | 3,4,5 | 3.8 | 7.5 |
| 22,033,993 | 1 | 684 | 4,5 | 1.9 | |
| 22,033,509 | 1 | 200 | 5 | 1.9 | |
| 22,033,508 | 4 | 199 | 5 | 7.5 | |
| 22,033,486 | 2 | 177 | 5 | 3.8 | |
| 22,033,485 | 2 | 176 | 5 | 3.8 | 1.9 |
| 22,033,473 | 11 | 164 | 5 | 20.8 | |
| 22,033,438 | 1 | 129 | coding | 1.9 | |
| 22,033,422 | 2 | 113 | coding | 3.8 | |
| 22,033,422 | 7 | 113 | coding | 13.2 | |
| 22,033,416 | 7 | 107 | coding | 13.2 | |
| 22,033,407 | 3 | 98 | coding | 5.7 | |
| 22,033,382 | 2 | 73 | coding | 3.8 | 37.7 |
| 22,033,377 | 1 | 68 | coding | 1.9 | 43.4 |

**Table S1 –** Summary of 5'RACE analysis showing the positions of 5 ' ends determined by 5'RACE sequencing, the number of clones retrieved for each end, the position of each 5'end in relation to the annotated human start site (Human version hg19- http://genome.ucsc.edu), the coverage or whether the sequence can be detected by the primers shown in fig S1D, and the percentages of clones determined for different coverage groups depicted in fig S1D.

**Movie S1.**

Time-lapse confocal microscopy (7 frames per second) of Vero E6 cells co-expressing (24 h) the Golgi marker Scarlet-Giantin, ZDHHC20$^{Long}$-RUSH-GFP reporter and an ER-resident hook. Synchronized trafficking was monitored upon D-biotin addition, after $T_0$ (see also Fig 4L and fig S4F). Scale bar 10 μm.

**Movie S2.**

Time-lapse confocal microscopy (7 frames per second) of Vero E6 cells co-expressing (24 h) the Golgi marker Scarlet-Giantin, ZDHHC20short-RUSH-GFP reporter and an ER-resident hook. Synchronized trafficking was monitored upon D-biotin addition after $T_0$ (see also Fig 4L and fig S4F). Scale bar 10 μm.
